# Unified multimodal learning enables generalized cellular response prediction to diverse perturbations

**DOI:** 10.1101/2025.11.13.688367

**Authors:** Chen Li, Lei Wei, Xuegong Zhang

## Abstract

Cells respond to diverse external interventions through shared regulatory mechanisms, suggesting that diverse interventions may be amenable to unified computational modeling. In practice, however, these responses are profiled across highly heterogeneous experimental settings, including distinct perturbation modalities, dosages, combinations, and cellular contexts. As a result, most computational models remain narrowly tailored to a single perturbation modality or experimental setting. They are difficult to extend to new cell types or perturbation types for which limited training data are available, and offer limited capacity to reuse information across heterogeneous perturbation datasets. Here, we introduce X-Pert, a general perturbation modeling framework that jointly represents external interventions and cellular contexts within a unified multimodal architecture. X-Pert adopts a mechanism-aligned design that explicitly models gene–perturbation interactions together with gene–gene dependencies through dedicated attention mechanisms, enabling the unified handling of heterogeneous experimental settings. Across benchmarks involving both genetic and chemical perturbations, X-Pert consistently outperforms existing methods under conventional accuracy metrics as well as biology-aware evaluations. Importantly, X-Pert exhibits strong generalization across cell types and supports joint learning across perturbation types. The unified latent space learned by X-Pert further enables downstream analyses such as perturbation retrieval and drug–gene association discovery, facilitating the prioritization of candidate gene inhibitors and the identification of anti-cancer compounds. Together, X-Pert establishes a versatile and generalizable foundation for perturbation modeling and predictive virtual cells, with broad applications in biomedical research and therapeutic discovery.

## Introduction

Cells respond to external interventions by transitioning between states^1^. These interventions can take diverse forms, ranging from genetic and chemical perturbations to combinatorial treatments, and may act on single molecular targets or across multiple components of the regulatory system^2^. Despite this diversity, such interventions reflect the same underlying process: an external operation applied to a cellular system that induces a biological response. This process is governed by gene regulatory mechanisms that are partially shared across perturbation types and cellular contexts^3^. Consequently, intervention response measurements obtained under different perturbations and cellular contexts can be viewed as complementary observations of common regulatory systems, motivating the development of general perturbation models that jointly capture heterogeneous perturbations across diverse cellular contexts within a unified framework, thereby enabling more efficient integration of heterogeneous perturbation data and supporting cross-perturbation knowledge transfer, with downstream implications for drug discovery and precision therapeutic strategies.

The rapid advancement of perturbation-based omics technologies^4-19^ has brought general perturbation modeling within practical reach. Perturbation-related cellular datasets now span a wide range of experimental settings encompassing bulk-level^20^ and single-cell^4^ measurements, diverse perturbation types such as genetic^4- 10^, chemical^11-15^, and protein^16-19^ perturbations, varying perturbation strengths including drug dosages^15^ and gene perturbation efficacies^21^, and a broad spectrum of cellular contexts across multiple cell types^7^. In parallel with these developments, perturbation datasets have expanded markedly in scale, from early studies profiling thousands of cells^4^ to modern resources containing tens of millions^15^. As perturbation data have grown in scale and diversity, perturbation modeling is beginning to move beyond collections of task-specific analyses toward general perturbation models that integrate heterogeneous data across contexts. When trained at scale within a unified framework, such models can leverage data diversity to learn increasingly rich regulatory representations, leading to qualitative improvements in generalization that are inaccessible to fragmented approaches.

A general perturbation model must simultaneously support unified and extensible modeling across perturbation types and robust generalization across cellular contexts. However, most existing computational approaches^22-35^ for *in silico* perturbation remain narrowly scoped and fragmented. The majority of current approaches are designed for a single perturbation type^22-25, 29-32^, typically focusing on either genetic perturbations^17-20^ or chemical perturbations^24-27^, and are often evaluated within a fixed cellular context. Indeed, most current approaches and evaluation settings, including those adopted in a recent community challenge^36^, focus on generalization within individual cell lines, which substantially limits the practical applicability of *in silico* perturbation models. While some methods, such as CellFlow^33^, aim to broaden perturbation modeling across different perturbation types, their scope remains limited to fixed cellular contexts, with evaluations largely confined to individual datasets. Conversely, approaches such as STATE^34^ emphasize generalization across cellular contexts, but remain limited in their ability to generalize across perturbation types. Taken together, these efforts advance complementary aspects of generalization, but remain insufficient for realizing general perturbation models that jointly integrate perturbation diversity and cellular context within a unified framework.

To address this gap, we propose X-Pert, a general perturbation modeling framework based on multimodal learning. X-Pert is motivated by the observation that external perturbations and cellular contexts represent distinct but interacting sources of information: perturbations specify external interventions applied to the system, while cellular contexts characterize the internal regulatory state that shapes how these interventions are realized. Accordingly, X-Pert models external perturbations and cellular states as distinct modalities, capturing perturbation–cell interactions as well as the propagation of regulatory effects through gene–gene dependencies that collectively shape cellular state transitions. By jointly modeling perturbation diversity and cellular context within a unified framework, X-Pert supports perturbation-agnostic prediction and robust generalization across cellular contexts. Together, these design principles establish X-Pert as a practical and extensible foundation for general perturbation modeling.

Using extensive benchmarks spanning genetic and chemical perturbations, we show that X-Pert consistently outperforms existing approaches under both conventional accuracy metrics and biology-aware evaluations that emphasize perturbation-specific and pathway-level responses. Across these comprehensive benchmarks, X-Pert consistently outperforms existing methods by achieving higher predictive accuracy and better capturing biologically meaningful variations among perturbations. X-Pert also generalizes across cell types, and its performance scales with cellular diversity in the training set, highlighting the benefits of learning from heterogeneous contexts at scale. Beyond single-modality tasks, X-Pert effectively harmonizes data from genetic and chemical perturbations to enable joint learning. By embedding perturbations with varying dosages, combinations, and types into a unified latent space, X-Pert facilitates diverse downstream applications, including genetic perturbation analysis, drug retrieval, and drug–gene association discovery. Collectively, these capabilities position X-Pert as a versatile and generalizable framework for *in silico* perturbation, offering a systematic foundation for understanding and designing cellular perturbations.

## Results

### X-Pert: a unified model for *in silico* perturbation

We designed X-Pert to predict the gene expression profile of a perturbed cell from the original cellular state and the corresponding intervention specification. Rather than framing perturbation response prediction as a collection of task-specific problems, X-Pert is formulated as a general perturbation model that treats external interventions and cellular contexts as distinct but interacting sources of information. Within a single unified architecture, X-Pert accommodates heterogeneous experimental settings, including bulk and single-cell measurements, diverse perturbation types such as genetic and chemical interventions, graded perturbation strengths (for example, drug dosage or gene perturbation efficacy), single and combinatorial perturbations, and diverse cellular contexts. By integrating perturbation representations and cellular states through a unified multimodal learning framework, X-Pert maps these heterogeneous inputs to a common predictive space, enabling consistent modeling of cellular responses across perturbation modalities and experimental conditions (Fig. 1A).

**Fig. 1.**
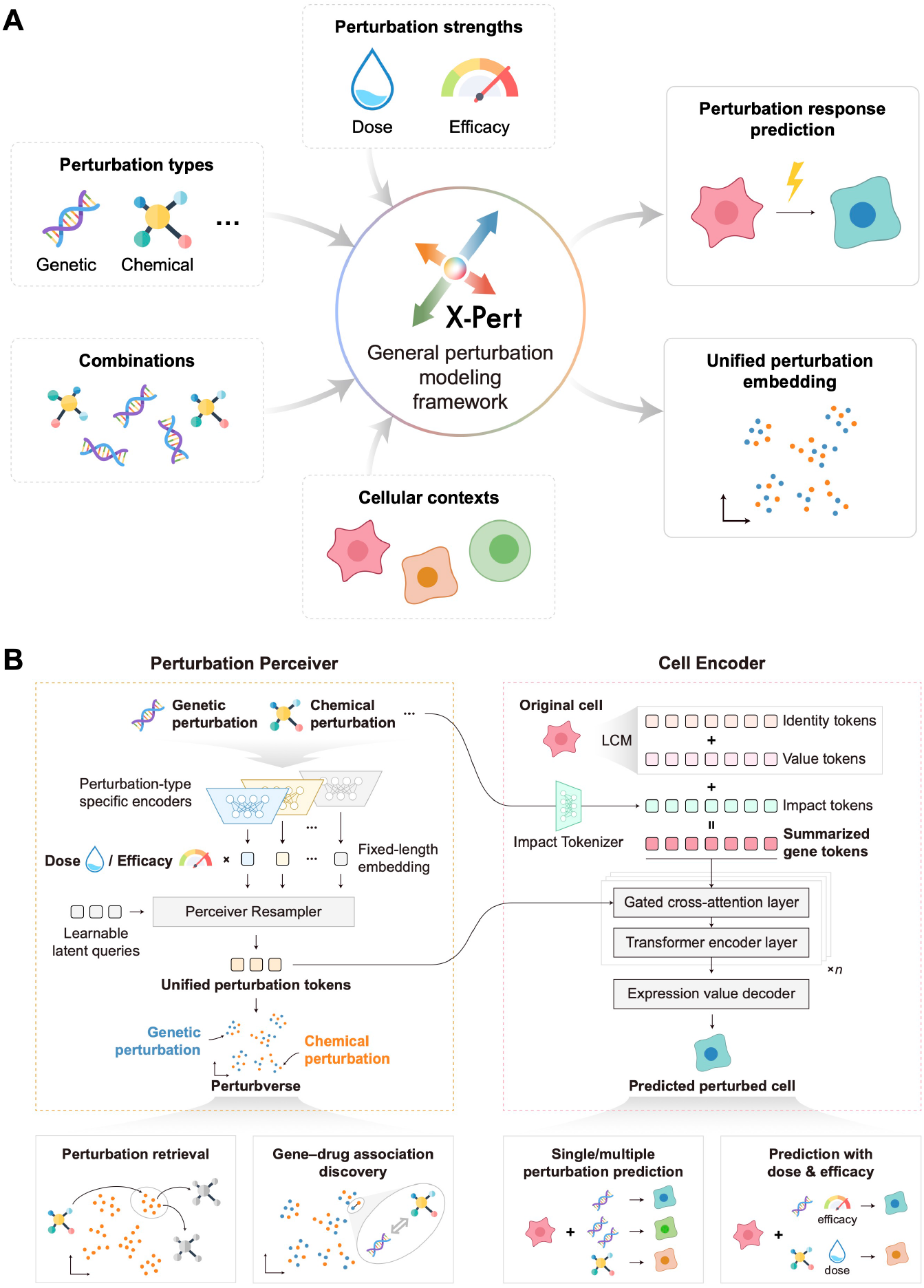
Overview of the model architecture of X-Pert. **(A)** X-Pert formulates perturbation response prediction as a general multimodal learning problem, integrating heterogeneous perturbation modalities (genetic and chemical), graded strengths (e.g., dosage or efficacy), single and combinatorial interventions, and diverse cellular contexts into a unified predictive framework. **(B)** X-Pert consists of two core components: the Perturbation Perceiver and the Cell Encoder. The Perturbation Perceiver is designed to accommodate different types of external perturbations, including multiple simultaneous perturbations and dose- or efficacy-dependent effects. Through a set of learnable latent queries, it projects perturbations of any type and number into a fixed-length latent representation, termed the Perturbverse. The Cell Encoder models the original cell using three types of tokens, namely identity tokens, value tokens, and impact tokens, which respectively encode gene identity, expression magnitude, and perturbation influence. By combining a gated cross-attention module with transformer encoder layers, the Cell Encoder integrates perturbation information with original gene expression to predict the transcriptional profile of perturbed cells. X-Pert can be applied to a wide range of perturbation-based tasks. Leveraging the Perturbverse, X-Pert enables perturbation retrieval and gene–drug association discovery through similarity analysis. Using the Cell Encoder’s predictive capability, X-Pert supports single/combinatorial perturbation prediction as well as modeling of perturbation efficacy and dose–response relationships.

The model consists of two key components: the Perturbation Perceiver and the Cell Encoder (Fig. 1B, Methods). The Perturbation Perceiver maps perturbations of different types into a shared latent representation space, providing a unified and extensible representation of external interventions that is independent of perturbation modality. Each perturbation type is first initialized as a fixed-length embedding by a type-specific encoder (in this work, we used text embeddings from GPT-4o^37^ for genetic perturbations, and SMILES-based embeddings computed with RDKit^38^ for chemical perturbations). Dose or perturbation efficacy information, when available, is then incorporated by augmenting the corresponding perturbation embedding. Depending on the experimental setting, a single perturbation embedding or a set of embeddings for multiple perturbations is then passed through a Perceiver Resampler, where a fixed set of learnable latent query tokens are employed to transform these embeddings into unified perturbation tokens (Methods). These tokens define a shared, length-invariant perturbation representation space that supports the unified modeling of heterogeneous perturbations and their combinations, which we term the Perturbverse.

The unified perturbation tokens produced by the Perturbation Perceiver serve as conditional context for the Cell Encoder, a transformer-based module designed to predict perturbed gene expression profiles by modeling both gene–perturbation interactions and gene–gene dependencies (Methods). The Cell Encoder builds upon a large cellular model (LCM) to encode an original cell as a sequence of gene identity tokens and expression value tokens (Methods). To better incorporate perturbation effects, we introduce an Impact Tokenizer that encodes perturbation-specific influences on each gene. The formulation of impact tokens is designed to flexibly accommodate different perturbation types: for genetic perturbations, they are modeled as one-hot vectors marking perturbed genes; for chemical perturbations, they are derived from protein–molecule affinity scores predicted by a pre-trained drug–target binding affinity prediction model^39^ (Methods). Each gene token is then summarized by integrating its gene identity, expression value, and impact components. These summarized gene tokens interact with perturbation tokens through cross-attention to capture perturbation– gene relationships to reflect how external interventions are realized through shared gene regulatory mechanisms. Subsequently, self-attention layers are employed to model gene–gene dependencies and preserve the intrinsic regulatory structure of the cell. The Cell Encoder outputs normalized gene expression values for the perturbed cell.

X-Pert is trained end-to-end with a mean squared error (MSE) loss between predicted and observed profiles (Methods). Once trained, it serves not only as a robust predictor of cellular responses but also as a versatile framework for downstream biological discovery. The Perturbverse serves as a unified reference space that enable cross-perturbation comparison and knowledge transfer, supporting perturbation retrieval and gene– drug association discovery. The Cell Encoder flexibly handles diverse datasets and biological contexts, supporting the prediction of single or multiple perturbations as well as dose- or efficacy-dependent responses. Together, these capabilities establish X-Pert as a practical realization of general perturbation modeling and provide a scalable foundation for downstream biological and therapeutic applications.

### Predicting responses of unseen genetic perturbations

We first evaluated X-Pert’s performance on unseen genetic perturbation prediction, representing a real-world use case in which the model infers cellular responses to perturbations not yet experimentally profiled within the same cellular context (Fig. 2A). To evaluate this capability, we benchmarked X-Pert on three single-cell perturbation datasets spanning CRISPR interference^7^ (CRISPRi, Replogle2022_K562 and Replogle2022_RPE1) and CRISPR activation^40^ (CRISPRa, Norman2019), comprising between 224 and 1,299 unique perturbations (Methods). For each dataset, perturbations were split 8:2 into training and testing (Methods). We compared X-Pert with six representative methods, including linear models^41^ (Linear, Linear-scGPT), deep learning methods (CellOracle^42^, GEARS^22^), and large cellular foundation models (scGPT^25^, scFoundation^24^), and the mean predictor (AveKnown) that have been used as a naive baseline in some studies^33, 43^.

**Fig. 2.**
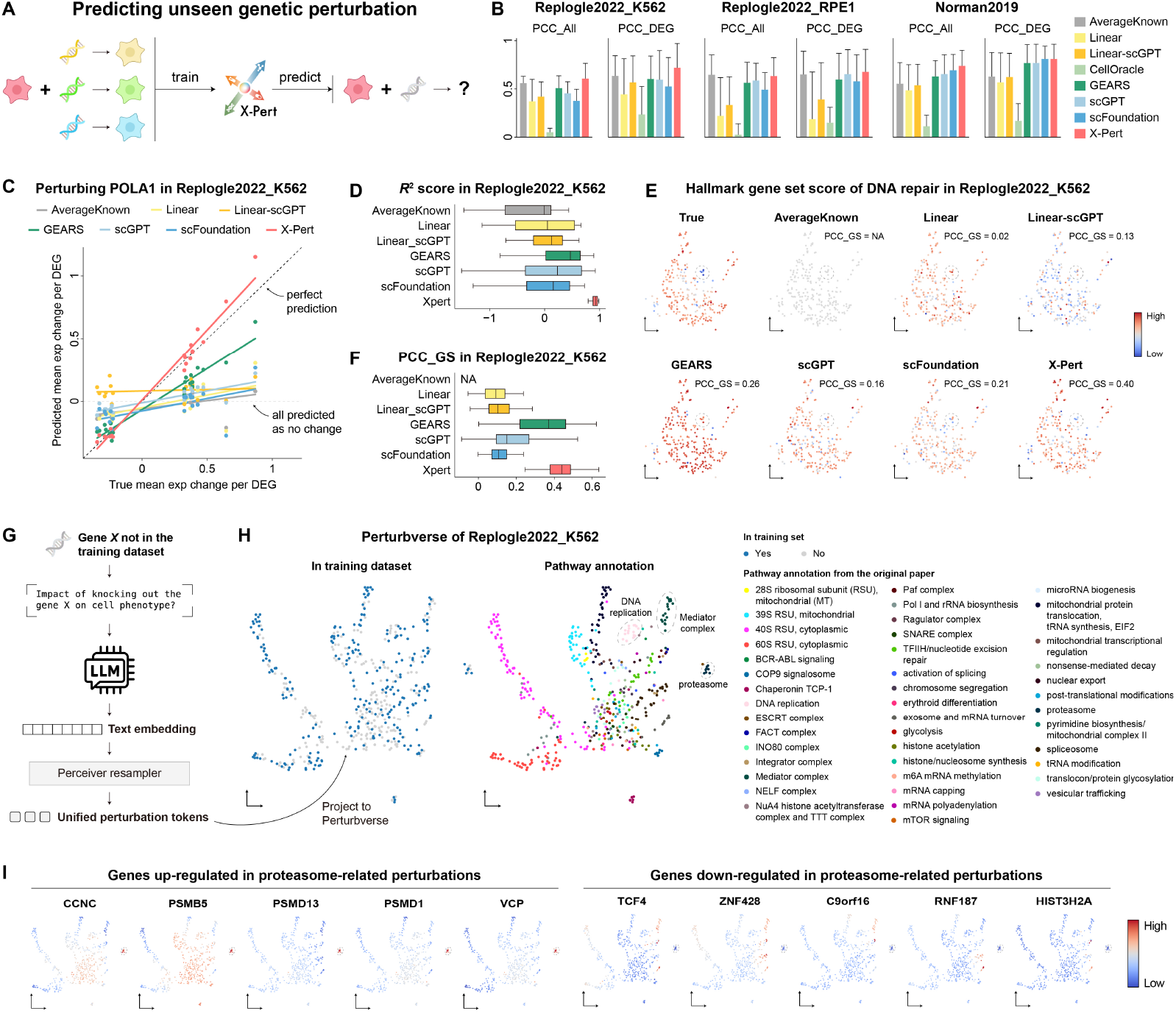
X-Pert’s predictions of genetic perturbations. **(A)** Schematic illustration of predictions on unseen genetic perturbations. **(B)** Comparison of X-Pert with other methods. **(C)** Comparison between predicted and ground-truth expression changes of the top 20 DEGs following POLA1 perturbation in Replogle2022_K562. Dots represent the mean predicted expression changes from each method, and a linear regression fit was added to visualize the correlation between the predicted and true mean values. CellOracle is not included because it only performs prediction on transcription factors. **(D)** Box plots comparing the *R*^2^ score between predicted and observed expression changes of top 20 DEGs in Replogle2022_K562. **(E)** UMAP visualization of Hallmark DNA repair gene set scores. The leftmost panel shows ground-truth scores, and each point represents a genetic perturbation. The PCC_GS between predicted and observed gene set scores is shown in each panel. The dashed circle highlights the ground-truth clustering of perturbations with reduced gene set scores. **(F)** Box plots comparing the PCC_GS of all 50 Hallmark gene sets across 260 test perturbations in Replogle2022_K562. For bar plots, bar: mean performance value; error bar: standard deviation across replicates. For box plots, center line, median; box limits, upper and lower quartiles; whiskers, 1.5× interquartile range. **(G)** Schematic illustration of using latent tokens from the Perturbverse to explore the functional landscape of genetic perturbations. **(H)** UMAP visualization of 504 genetic perturbations from Replogle2022_K562, with pathway annotations manually curated from the original dataset. **(I)** UMAP visualization of mean gene expressions across 504 annotated genetic perturbations from Replogle2022_K562. The ten displayed genes were selected from significant DEGs in proteasome-related perturbations. Dashed circles represent cells perturbed by proteasome-related genes.

We first evaluated model performance using a commonly used metric, the Pearson correlation coefficient (PCC) across all genes, referred to as PCC_All, which quantifies the correlation between predicted and observed gene-expression changes for each perturbation, calculated over all genes as paired values of predicted and measured expression differences relative to the original cells. As shown in Fig. 2B, X-Pert achieved the best or highly competitive performance on PCC_All across all three datasets. However, we found that PCC_All provides little meaningful differentiation among models, as even naïve predictors such as AveKnown perform similarly. A recent study has revealed a pervasive systematic bias in genetic-perturbation datasets, where shared global response components dominate gene-expression variance, overshadowing the true perturbation-specific regulatory signals^44^. Consequently, even simple baselines such as AveKnown can achieve deceptively high correlation scores by reproducing these shared patterns, a phenomenon consistently reported across benchmarks^41, 43^. Therefore, while PCC_All remains a convenient and widely used summary statistic, it primarily reflects the recovery of background structure rather than the accurate modeling of biological perturbation effects.

To better assess biological relevance responses, we next examined PCC_DEG, which quantifies the correlation between predicted and observed expression changes only among DEGs (Methods). This metric filters out nonspecific or noisy variation and provides a clearer view of biologically meaningful transcriptional responses. For instance, it reached 0.720 on Replogle2022_K562, a 13.2% improvement over the second-best method, with similar gains observed on Replogle2022_RPE1 (+3.8%). Importantly, most methods showed a larger increase in performance when evaluated by PCC_DEG compared with PCC_All across datasets. This consistent widening of the performance gap suggests that PCC_DEG captures perturbation effects more faithfully by highlighting genuine expression changes associated with each perturbation, which are largely averaged out in PCC_All by shared global responses. However, despite this improvement, the PCC_DEG of AveKnown remains high, indicating that this metric still carries dataset-level correlations and is limited in distinguishing biologically meaningful variation from background effects. These limitations underscore the need for evaluation strategies that move beyond correlation-based similarity and instead assess a model’s ability to reconstruct both the magnitude and the functional organization of true perturbation effects.

To move beyond correlation-based metrics and directly assess how accurately each model captures real perturbation effects, we focused on perturbations with at least 20 significant DEGs. This subset reflects conditions where transcriptional regulation is strongly altered and thus provides a clearer test of biological fidelity. For each perturbation, we compared the predicted and observed expression changes of the top 20 DEGs, which represent the most strongly responsive genes and are typically related to the downstream functional consequences of the perturbation. This evaluation highlights not only whether a model predicts the correct direction of change but also whether it reproduces the actual magnitude of transcriptional shifts. For example, in the POLA1 perturbation of Replogle2022_K562, X-Pert closely recapitulated both the direction of magnitude of top 20 DEGs’ expression changes, whereas other methods tended to compress predictions toward low-magnitude shifts (Fig. 2C). Similar patterns were observed for additional perturbations across datasets (Fig. S1A). We further quantified this agreement using the coefficient of determination (*R*^2^) between the predicted and observed expression changes of the top DEGs (Methods). Across all three datasets, X-Pert consistently achieved the highest *R*^2^ scores (Fig. 2D and Fig. S1B), demonstrating its ability to accurately reproduce the real magnitudes of perturbation-induced transcriptional responses rather than merely capturing overall expression trends.

While single-gene evaluations help assess whether a model can reproduce individual expression changes, cellular responses are in fact mediated by coordinated regulation of gene programs. Therefore, evaluating predictions at the pathway level provides a more biologically meaningful and noise-robust assessment of model performance. To assess pathway-level responses, we computed Hallmark gene set scores^45^ for each perturbation by averaging the corresponding signature scores across all cells under that condition (Methods), reflecting the activation state of specific pathways after perturbation. Such evaluation moves beyond conventional correlation-based metrics, as it directly tests whether a model can recover the coordinated activation or repression of biological programs that define a cell’s functional state. We quantified the agreement between predicted and observed gene set scores across different perturbations using PCC (PCC_GS). Unlike gene-level metrics that assess agreement within each perturbation, PCC_GS compares model performance across perturbations, evaluating whether the model can correctly rank perturbations by the magnitude and direction of their pathway-level effects. This distinction is important because the ability to prioritize perturbations by their functional impact forms the basis for many biological applications such as perturbation retrieval, target identification, and mechanism inference.

On 260 test perturbations from the Replogle2022_K562 dataset, X-Pert captured this diversity with high fidelity. For example, in the DNA repair gene set, it achieved the highest PCC_GS (0.40), representing a 53.8% improvement over the second-best method, GEARS (PCC_GS = 0.26, Fig. 2E). In contrast, AveKnown completely failed under this setting, producing identical predictions for all perturbations and thus showing no ability to distinguish functional differences between them. Furthermore, X-Pert successfully predicted a cluster of perturbations exhibiting coordinated decreases in DNA repair gene set scores within the UMAP space, a structure that none of the other methods were able to capture (Fig. 2E). Similar advantages were observed for additional gene sets such as adipogenesis, apical junction, and protein secretion (Fig. S1C). Across all 50 Hallmark gene sets, X-Pert consistently outperformed all other methods (Fig. 2F), demonstrating superior ability to model perturbation-specific gene program changes.

These results demonstrate that X-Pert not only achieves accurate performance across multiple evaluation metrics but also provides biologically coherent predictions at both gene and pathway levels. By evaluating relative perturbation effects across perturbations rather than isolated gene-level accuracy within individual perturbations, pathway-level benchmarks provide a more faithful assessment of general perturbation modeling performance, consistent with recent perturbation benchmarks^44^ that emphasized evaluating performance across the full spectrum of perturbations rather than isolated cases.

Building on X-Pert’s strong predictive performance, we next examined what the model has learned about the underlying structure of perturbations. We explored the interpretability of X-Pert’s learned latent perturbation space, or Perturbverse, generated by the Perturbation Perceiver (Methods). Because perturbations are represented using gene-related text embeddings, even perturbations absent from training can be directly embedded into the space (Fig. 2G). This provides a structured representation of perturbation effects for revealing functional relationships among genetic perturbations. For instance, 504 genetic perturbations from Replogle2022_K562 revealed clusters corresponding to biological processes such as proteasome, mediator complex, and DNA replication, despite only 209 perturbations being included in training (Fig. 2H and S2A).

The Perturbverse also provides a convenient framework to examine how clusters of perturbation genes relate to downstream expression changes. By mapping all input perturbations into the Perturbverse (Methods), we found that proteasome-related perturbations formed a coherent cluster, and their predicted profiles showed coordinated upregulation of multiple subunits (PSMB5, PSMD1, PSMD13), consistent with the known “bounce-back” transcriptional response^46, 47^ (Fig. 2H–I). A similar Perturbverse constructed for Replogle2022_RPE1, despite being trained on only 73 perturbations, also recovered clear and biologically coherent structures (Fig. S2A–B). These results highlight that X-Pert not only achieves accurate prediction of unseen genetic perturbations but also organizes them into an interpretable latent space that reveals functional relationships between perturbation clusters and transcriptional programs.

### Predicting responses under perturbation combinations and considering efficacy

The flexible design of the Perturbation Perceiver enables X-Pert to naturally handle both combinatorial perturbations and perturbation efficacy. To assess its ability to generalize to unseen combinations (Fig. 3A), we used the Norman2019 dataset, which includes 148 single-gene and 76 two-gene CRISPRa perturbations. To simulate the unseen-combination setting, we randomly excluded 30% of the single-gene perturbations during training and evaluated two types of double perturbations: one-unseen combinations (*n* = 61), where one perturbed gene was absent from training, and two-unseen combinations (*n* = 15), where both were unseen. Notably, the evaluation did not include “zero-unseen” combinations, as such cases do not naturally arise from this data-splitting strategy.

**Fig. 3.**
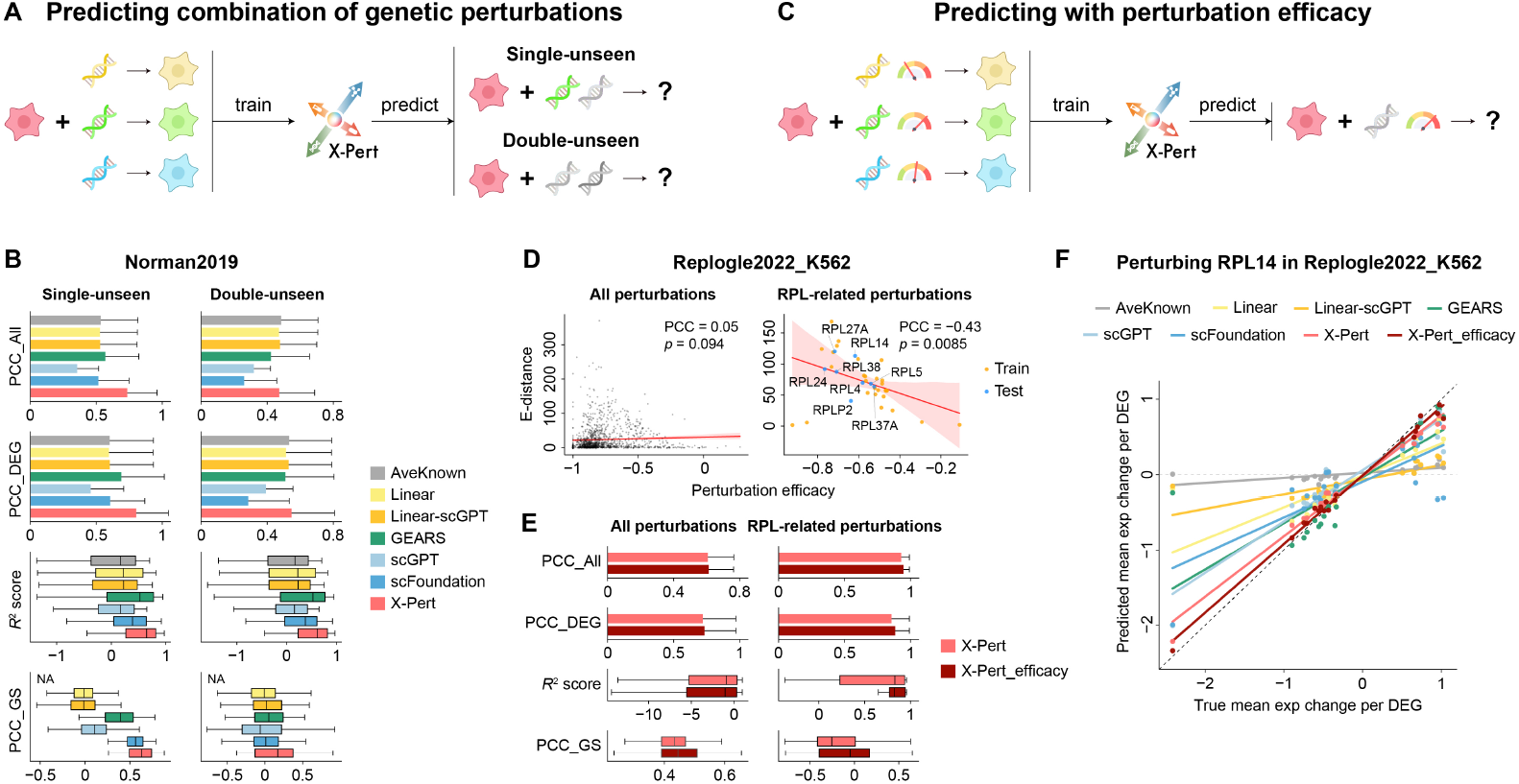
X-Pert’s predictions of perturbation combinations and with efficacy modeling. **(A)** Schematic illustration of multi-perturbation prediction. **(B)** Comparison of X-Pert with other methods in the combinatorial perturbation prediction setting. For bar plots, bar: mean performance value; error bar: standard deviation across replicates. For box plots, center line, median; box limits, upper and lower quartiles; whiskers, 1.5× interquartile range. **(C)** Schematic illustration of perturbation prediction incorporating efficacy information. **(D)** Scatter plots with fitted regression lines showing the relationship between E-distance and perturbation efficacy across all perturbations (left) and RPL-related perturbations (right) in Replogle2022_K562. **(E)** Bar plots comparing X-Pert and X-Pert_efficacy across all test perturbations (left) and RPL-related perturbations (right). **(F)** Comparison between predicted and ground-truth expression changes of the top 20 DEGs following RPL14 perturbation in Replogle2022_K562. Dots represent the mean predicted expression changes from each method, and a linear regression fit was added to visualize the correlation between the predicted and true mean values.

As shown in Fig. 3B, X-Pert substantially outperformed all other methods in the one-unseen setting across all metrics. Even in the more challenging two-unseen setting, X-Pert maintained the best performance in in *R*^2^ score and PCC_GS, and remained highly competitive in PCC_All and PCC_DEG. Notably, X-Pert was the only model that achieved a clearly positive median PCC_GS in the two-unseen setting, underscoring its capacity to generalize to novel perturbations by leveraging gene-related text embeddings for perturbation representation. Together, these results demonstrate that X-Pert effectively integrates multiple perturbations and extends robustly to unseen combinatorial conditions.

Modeling perturbation efficacy represents another important challenge (Fig. 3C), as perturbation strength is a critical but often under-modeled attribute of external interventions. For example, In CRISPR-based experiments, perturbation efficacy depends heavily on sgRNA design but is not directly measurable, unlike chemical perturbations where dose provides a natural quantitative proxy. To address this, we quantified genetic perturbation efficacy by measuring target gene downregulation relative to original cells (Methods), and examined its relationship with the overall transcriptional impact, quantified by the energy distance (E-distance) between perturbed and original profiles. For the full Replogle2022_K562 dataset, we didn’t find global correlation between E-distance and perturbation efficacy (Fig. 3D). This is likely because many perturbations induce weak or noisy transcriptional shifts, which obscure the link between efficacy and downstream effects. However, focusing on a more homogeneous subset of 37 ribosomal protein large subunit (RPL) perturbations, we observed a significant negative correlation (PCC = –0.43, *p* = 0.0085; Fig. 3D), indicating that higher efficacy generally drives stronger transcriptional responses. These results suggest that perturbation efficacy constitutes an important latent variable that should be explicitly incorporated into general perturbation models.

Building on this insight, we extended X-Pert to incorporate efficacy information, yielding X-Pert_efficacy (Methods). Across all perturbations in Replogle2022_K562, X-Pert_efficacy performed comparably to or slightly better than the original model (Fig. 3E). When investigating the RPL subset, it showed a clear advantage: X-Pert_efficacy achieved significantly higher *R*^2^ on eight held-out perturbations (Fig. 3E) and more faithfully recovered expression changes of top DE genes (Fig. 3F and Fig. S3). For example, in the case of RPL14 perturbation, it can more accurately predicted expression changes in top DEGs, which were underestimated by the original model (Fig. 3F). Similar improvements were consistently observed across other test RPL perturbations (Fig. S3), demonstrating that explicitly modeling perturbation efficacy improves biological fidelity and supports more accurate generalization across interventions of varying strength.

### Predicting responses of chemical perturbations

We next evaluated X-Pert on chemical perturbations (Fig. 4A), which provide a complementary and more complex class of interventions compared with genetic perturbations. We first benchmarked it on the Sciplex-3 dataset^14^, which profiles three cell lines (A549, K562, MCF7) treated with 188 drugs across four concentrations (10 nM–10,000 nM, 758,368 cells after preprocessing). Perturbations were split 8:2 into training and testing. We selected representative chemical perturbation prediction methods for comparison, including ChemCPA^29^, BioLord^30^, and PRnet^31^. As shown in Fig. 4B, X-Pert achieved the best or highly competitive performance across PCC_All, PCC_DEG, and *R*^2^ score, and reached the highest PCC_GS in K562 and MCF7. However, the overall predictive performance across all models was modest, with median *R*^2^ scores close to zero. This limitation appears to arise from the intrinsic characteristics of the Sciplex-3 dataset: most perturbations induced only a small number of significant DEGs, even with higher compound concentrations (Fig. S4A). Such weak transcriptional responses result in a low signal-to-noise ratio, making it difficult for models to learn robust perturbation-response relationships.

**Fig. 4.**
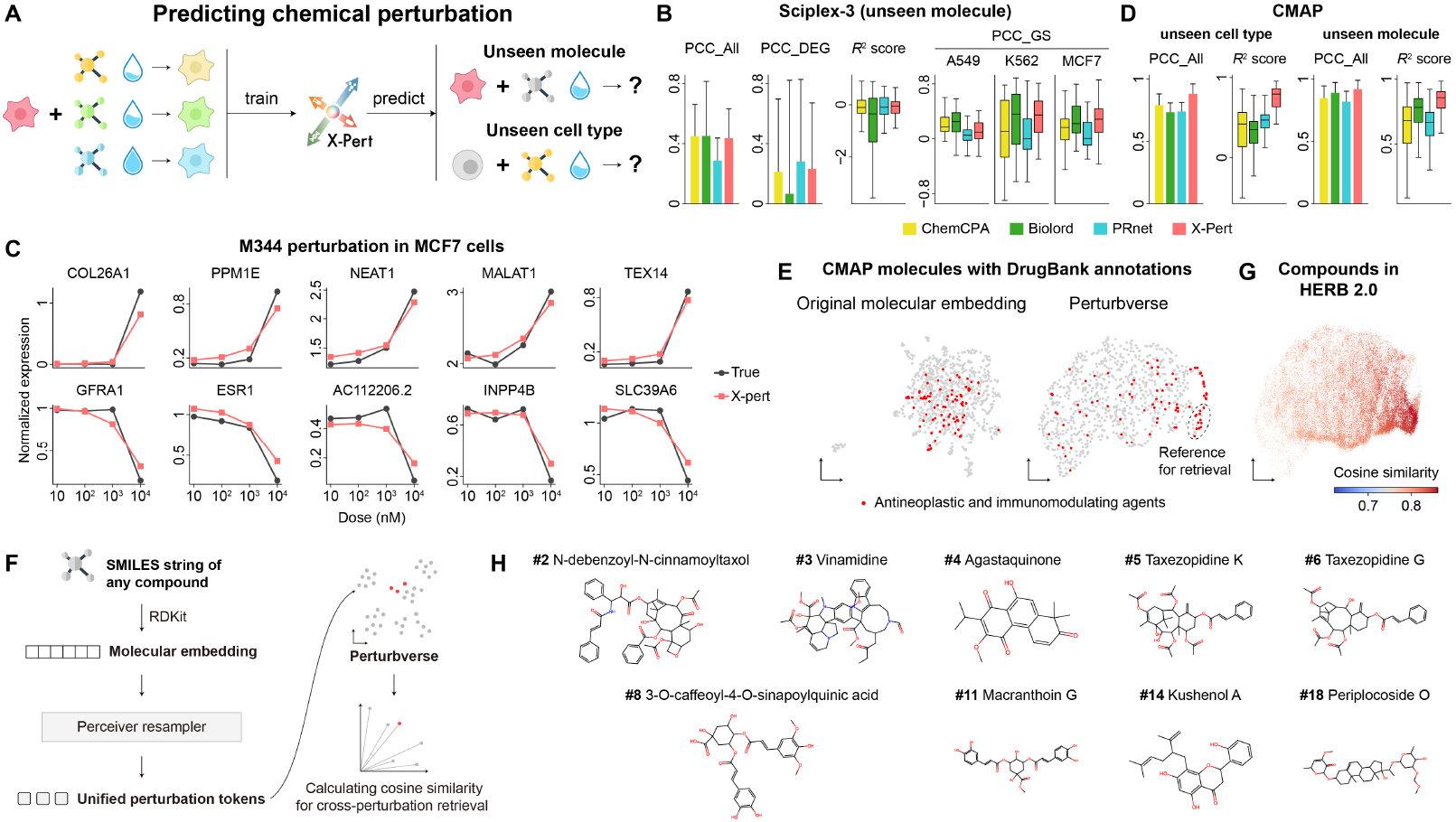
X-Pert predictions on chemical perturbations. **(A)** Schematic illustration of chemical perturbation prediction. **(B)** Model comparison on the Sciplex-3 dataset under the unseen molecule prediction scenario. **(C)** Line plots showing X-Pert–predicted gene expression levels versus ground-truth measurements across four chemical doses. **(D)** Model comparison on the CMAP dataset under both unseen molecule and unseen cell type settings. **(E)** UMAP visualization of chemical perturbations in the original molecular embedding space and the Perturbverse, colored by ATC Level 1 annotations. **(F)** Schematic illustration of the drug retrieval process using latent chemical embeddings derived from the Perturbverse. **(G)** UMAP visualization showing the cosine similarity distribution of candidate molecules from the HERB 2.0 database. **(H)** Top-ranked molecules retrieved by X-Pert from the HERB 2.0 database, shown with their rankings and compound names. For bar plots, bar: mean performance value; error bar: standard deviation across replicates. For box plots, center line, median; box limits, upper and lower quartiles; whiskers, 1.5× interquartile range.

To test whether X-Pert captures dose-dependent effects, we examined the M344 perturbation in MCF7 cells as a case study. M344 is a histone deacetylase (HDAC) inhibitor with marked dose-dependent heterogeneity, currently under investigation for its therapeutic potential in cancer and Alzheimer’s disease^48^. Across the top 10 DEGs, X-Pert faithfully reproduced dose–response patterns, capturing both trends and magnitudes of change (Fig. 4C). Similar results were observed across other compounds (Fig. S4B), confirming the effectiveness of explicit dose modeling within the unified perturbation framework.

We further extended the evaluation to the CMAP (Connectivity Map) dataset^20^, a large bulk-level perturbation resource profiling 17,775 small molecules across 70 cell lines using the L1000 technology (678,401 profiles after preprocessing). The scale and diversity of CMAP enable stringent evaluation of cross-molecule and cross-cell-type generalization, posing greater challenges than single-cell datasets. To account for bulk-level measurements, we evaluated both PCC_All and *R*^2^ (PCC_DEG and PCC_GS were excluded from the evaluation because the L1000 technology already selected 978 genes for profiling; *R*^2^ was calculated based on all profiled genes). As shown in Fig. 4D, X-Pert consistently outperformed all other methods under both settings. On unseen molecules, it exceeded BioLord by 3.1% in PCC_All and 12.0% in *R*^2^. On unseen cell types, the gains were even larger (PCC_All +11.4%, *R*^2^ +74.3%), highlighting X-Pert’s ability to generalize chemical perturbation responses across diverse cellular contexts.

We then explored whether chemical perturbations form meaningful representations in the Perturbverse. For 851 CMAP molecules with DrugBank ATC level 1 annotations^49^, original embeddings lacked discernible organization, whereas X-Pert grouped molecules into functionally coherent clusters, such as “antineoplastic and immunomodulating agents” (Fig. 4E and Fig. S4C). This organization emerges naturally from unified perturbation modeling, and is biologically plausible given that most CMAP profiles were generated from cancer-derived cell lines, likely amplifying signals from antineoplastic compounds.

Finally, we tested the utility of X-Pert for drug retrieval (Fig. 4F) by projecting ∼30,000 natural compounds from the natural compound database HERB 2.0^50^ into the Perturbverse and ranking them by cosine similarity to reference antineoplastic drugs (Fig. 4G, Methods). Among the top 20 retrieved candidates, nearly half have documented anticancer activity (Fig. 4H), including diterpenoid quinones^51^ (e.g., Agastaquinone), taxane-type diterpenoids^52, 53^ (Taxezopidines K and G), and caffeoylquinic acid derivatives^54, 55^ (3-O-caffeoyl-4-O-sinapoylquinic acid), all consistent with known pharmacological properties. In addition, structurally related analogues of paclitaxel (e.g., N-debenzoyl-N-cinnamoyltaxol) emerged, suggesting potential therapeutic relevance.

Together, these results demonstrate that X-Pert not only predicts chemical perturbation responses with high accuracy, including dose-dependent effects, but also organizes chemical perturbations into a unified and biologically meaningful latent space that supports cross-context generalization and enables *in silico* drug retrieval and discovery within a general perturbation modeling framework.

### *In silico* drug screening using large-scale chemical perturbation data

To extend X-Pert to large-scale *in silico* drug screening, we trained it on Tahoe-100M^15^, the largest publicly available single-cell chemical perturbation dataset to date. Generated with the Mosaic platform, Tahoe-100M comprises ∼100 million cells across 379 compounds, each profiled at three doses in 50 cell lines. The scale and diversity of this dataset provide a stringent setting for evaluating whether a unified perturbation architecture can effectively integrate perturbation signals across compounds, doses, and cellular contexts. To improve computational efficiency, we aggregated single-cell profiles into pseudo-bulk measurements (Methods), which preserved perturbation signals while enabling tractable training. Perturbations were split 8:2 into training and testing.

X-Pert accurately reconstructed the transcriptomic distributions of different cell lines in the UMAP space, clearly distinguishing them from one another (Fig. 5A), indicating that X-Pert captures cell line–specific regulatory contexts while operating at large scale. However, closer inspection of A549 cells revealed that plate labels introduced a strong batch effect, which masked drug-specific signals (Fig. 5B). To mitigate this, we incorporated a linear regression-based adjustment to remove batch-associated variation before drug screening (Methods). This correction ensured that downstream analyses reflected perturbation-induced biological effects rather than technical artifacts. After correction, we implemented an *in silico* drug screening pipeline (Fig. 5C, Methods): for each disease, we curated up- and down-regulated gene sets as disease signatures, computed gene set scores under candidate perturbations (denoted as up-scores and down-scores), and identified drugs with lower up-scores and higher down-scores with more potential that could reverse disease signatures. This strategy exploits X-Pert’s ability to model coordinated pathway-level responses to identify compounds whose predicted effects align with disease-relevant transcriptional programs.

**Fig. 5.**
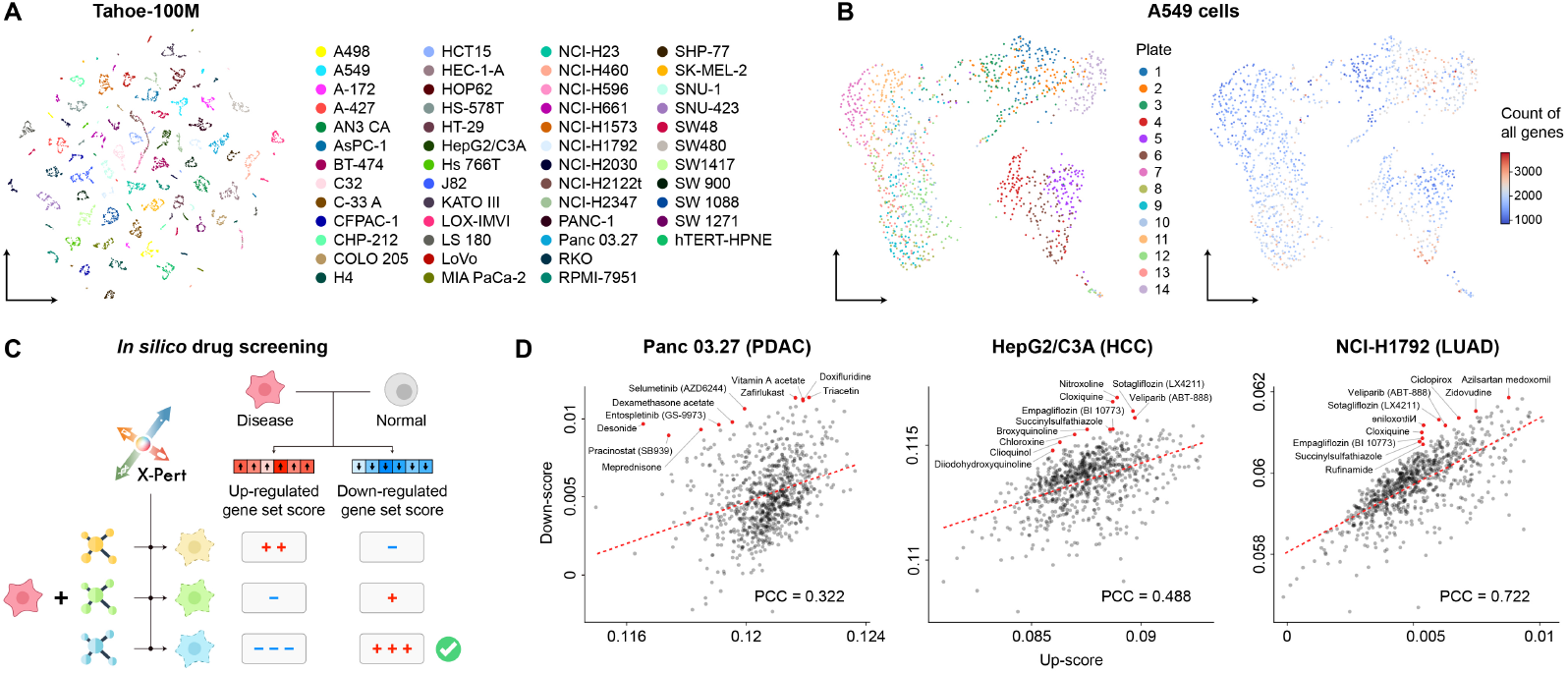
*In silico* drug screening using the Tahoe-100M dataset. **(A)** UMAP visualization of test perturbations predicted by X-Pert, where each point represents a pseudo-bulk cell derived from a specific chemical perturbation in a cell line. Points are colored by cell line. **(B)** UMAP visualization of all A549 samples, colored by sequencing plate and count of all genes, respectively, highlighting potential batch effects. **(C)** Schematic illustration of the *in silico* drug screening pipeline. **(D)** Scatter plots showing the top-ranked candidate drugs and fitted regression lines for three diseases: PDAC, HCC, and LUAD. Lower up-scores and higher down-scores represent drugs with higher potential that could reverse disease signatures. The top ten prioritized drugs are annotated in red.

We applied this approach to prioritize drugs for three cancer types: pancreatic ductal adenocarcinoma (PDAC; Panc 03.27), hepatocellular carcinoma (HCC; HepG2), and lung adenocarcinoma (LUAD; NCI-H1792). From an FDA-approved drug list, we excluded compounds used for training, yielding 831 candidates. X-Pert ranked the top ten drugs for each cancer.

As shown in Fig. 5D, for PDAC, top candidates included glucocorticoids (desonide, meprednisone, and dexamethasone acetate), which have shown unexpected anti-tumor activity in PDAC models^56^. X-Pert also highlighted Pracinostat, a pan-HDAC inhibitor under clinical investigation^57^. Another strong candidate was Entospletinib, a selective SYK inhibitor originally developed for B-cell cancers^58^, with evidence of reshaping tumor immunity^59^ and overcoming PDAC chemoresistance^60^, as supported by related compounds in preclinical models. For HCC, X-Pert identified nitroxoline, a cathepsin B inhibitor with well-documented anti-liver cancer effects^61^. Another candidate, clioquinol, an 8-hydroxyquinoline derivative, exerts anticancer activity through metal chelation, proteasome inhibition, and apoptosis induction^62^, and preclinical studies reported its antiproliferative effects in liver cancer cells^63^. Empagliflozin, an SGLT2 inhibitor approved for diabetes^64^ with emerging evidence of benefit in preclinical HCC models^65^, was also prioritized. For LUAD, X-Pert prioritized sotagliflozin, consistent with the reliance of LUAD on SGLT2-mediated glucose uptake^66^ and clinical signals of benefit from SGLT2 inhibitors in NSCLC patients^67^. It also highlighted ciclopirox, which suppresses proliferation and induces apoptosis in LUAD models^68^, and veliparib, a PARP inhibitor with reported benefit in NSCLC subgroups^69^.

Together, these results demonstrate that X-Pert can scale to massive perturbation datasets and integrate chemical perturbation responses with disease-specific transcriptional signatures, enabling systematic prioritization of candidate drugs across multiple cancer types. This large-scale in silico screening experiment illustrates how a general perturbation modeling framework can translate predictive modeling into actionable therapeutic hypotheses, supporting drug discovery and repurposing at scale.

### Performance scaling of X-Pert with increasing cell-type diversity

A defining challenge for general perturbation modeling is whether a model can benefit from perturbation data collected in cellular contexts different from the one of interest. While perturbation responses are known to be highly context-dependent, it remains unclear whether perturbation data from heterogeneous cell types can be systematically leveraged to improve prediction in a target cell type, or whether such heterogeneity instead introduces noise that limits generalization. To address this question, we investigated how X-Pert scales with increasing cellular diversity in the training data.

We assembled genome-wide CRISPRi perturbation datasets from four human cell lines profiled across two studies: Replogle2022_K562^7^, Replogle2022_RPE1^7^, Nadig2025_HepG2^70^, and Nadig2025_Jurkat^70^. These cell lines span diverse lineages, including K562 (a chronic myelogenous leukemia cell line derived from hematopoietic progenitors), RPE1 (an immortalized retinal pigment epithelial cell line), HepG2 (a hepatocellular carcinoma cell line), and Jurkat (a T-cell acute lymphoblastic leukemia cell line). To ensure consistency across cell types, we restricted the analysis to the 284 genetic perturbations shared across all four datasets, yielding a total of 255,194 cells. We designated K562 as the target cell type and split its perturbations into training and testing sets at a 3:7 ratio. All perturbation data from the remaining three cell lines were used exclusively as incremental training data and were incrementally incorporated during model training (Fig. 6A).

**Fig. 6.**
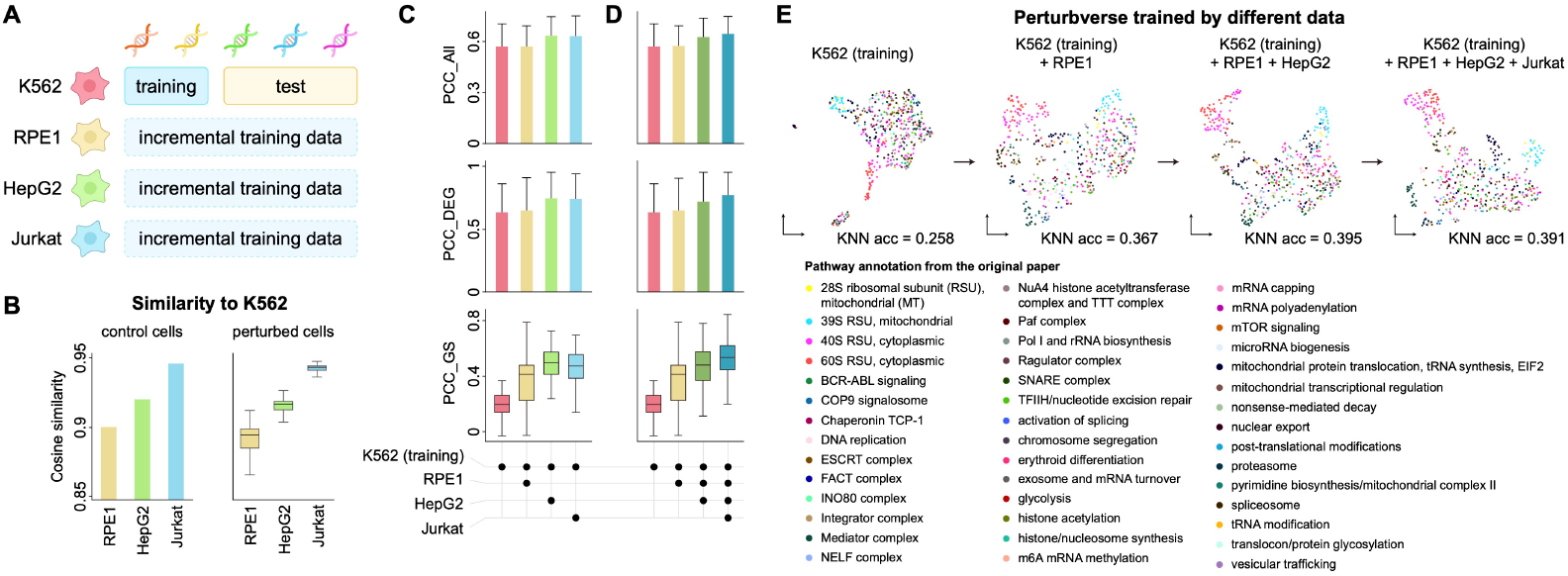
X-Pert scales with increasing cellular diversity in training data. **(A)** Schematic of the experimental design. K562 was designated as the target cell type, with its perturbations split into training and testing sets (3:7). Perturbation data from RPE1, HepG2, and Jurkat were used exclusively as incremental training data and were progressively incorporated during model training. **(B)** Pairwise cosine similarity of gene expression profiles across the four cell lines, computed for control cells and for perturbation-induced expression changes. **(C)** Predictive performance in K562 when incorporating perturbation data from a single external cell type. **(D)** Predictive performance as perturbation data from multiple external cell types are incrementally added. **(E)** Visualization of perturbation embeddings for 504 annotated human genes in the Perturbverse as additional cell types are incorporated.

We first examined the baseline similarity among these four cell types. Jurkat exhibited the highest similarity to K562 in control cells (cosine similarity = 0.946), consistent with their shared hematopoietic origin, whereas RPE1 showed the lowest similarity, reflecting its epithelial and non-cancerous nature (Fig. 6B). A similar trend was observed when comparing perturbation-induced expression changes across cell types (Fig. 6B), indicating that cellular relatedness partially shapes perturbation response similarity. These observations provided a biologically grounded expectation that perturbation data from some cell types may be more informative for K562 than others.

We then examined whether incorporating perturbation data from other cell types could indeed enhance predictive performance in K562. Interestingly, incorporating perturbation data from external cell types consistently improved predictive performance in K562 across all evaluation metrics (Fig. 6C). Even the addition of perturbation data from a single external cell type led to substantial gains, demonstrating that X-Pert can effectively transfer perturbation knowledge across cellular contexts. For example, PCC_GS increased from 0.186 when training on K562 alone to 0.372, 0.481, and 0.461 after incorporating data from RPE1, HepG2, or Jurkat, corresponding to relative improvements of 100.0%, 158.6%, and 147.9%, respectively. Similar improvements were observed for PCC_All and PCC_DEG. Notably, the magnitude of improvement correlated with biological similarity between cell types, with HepG2 and Jurkat yielding larger gains than RPE1, suggesting that cross–cell-type generalization reflects a combination of architectural scalability and biological relatedness.

We next asked whether these benefits continue to accumulate as perturbation data from additional cell types are incorporated. To address this question, we designed an experimental workflow in which perturbation data from different cell types were added incrementally during training, progressively adding perturbation data from RPE1, HepG2, and Jurkat to the K562 training set. We found that, across all metrics, performance improved monotonically with increasing cellular diversity (Fig. 6D). For instance, PCC_DEG increased from 0.640 when training on K562 alone to 0.779 when all four cell types were included, while PCC_GS improved from 0.186 to 0.517. These results reveal a clear scaling behavior, in which expanding cellular diversity in the training data leads to progressively improved generalization rather than diminishing returns.

To investigate how cross–cell-type scaling affects representation learning, we analyzed the evolution of genetic perturbation embeddings in the Perturbverse as additional cell types were incorporated. Following the procedure used in Fig. 2H, we visualized perturbation embeddings of 504 human-annotated genes. When X-Pert was trained solely on K562 data, the learned gene embeddings were poorly organized, with limited alignment to known gene annotations and a low KNN accuracy of 0.258 (Fig. 6E, Methods). As perturbation data from additional cell types were added, the embedding space became increasingly structured and biologically coherent, with KNN accuracy rising to 0.367, 0.395, and 0.391 (Fig. 6F). This progressive organization of the Perturbverse indicates that cellular diversity enriches the learned representation of gene function.

Together, these results demonstrate that X-Pert systematically benefits from perturbation data spanning diverse cellular contexts. Rather than degrading performance, cellular heterogeneity provides a source of transferable regulatory information that improves both predictive accuracy and representation quality. This scaling behavior supports the view that general perturbation models can accumulate knowledge across cell types within a unified framework, reinforcing X-Pert’s role as a scalable foundation for perturbation modeling across heterogeneous biological contexts.

### Aligning genetic and chemical perturbations in Perturbverse

The unified design of the Perturbation Perceiver allows X-Pert to jointly model multiple perturbation types within a unified framework, enabling direct comparison and functional alignment across distinct perturbation types. To evaluate this capability, we used the CMAP dataset, which contains both genetic and chemical perturbations profiled on same cell lines. We focused on three cell lines with the largest number of dual– perturbation-type profiles: A549 (21,685 chemical and 3,620 genetic perturbations), VCAP (34,753 chemical, 3,641 genetic), and HT29 (14,513 chemical, 3,302 genetic), and trained X-Pert independently on each cell line with an additional Maximum Mean Discrepancy (MMD) loss to encourage cross-type alignment (Methods).

We first quantitatively evaluated X-Pert’s ability to align genetic and chemical perturbations. Visualization of the Perturbverse showed that genetic and chemical perturbations were well integrated (Fig. 7A), rather than forming type-specific clusters. We selected a set of representative genes, including AKT1, AKT2, CTSK, SMAD3, TP53, MTOR, BRAF, and PIK3CA, as they represent diverse, therapeutically relevant signaling pathways and are associated with known or putative small-molecule modulators. For each gene, we queried GDIdb^71^ to identify drugs interacting with it, yielding 7–23 compounds per gene. We then calculated the cosine similarity between the latent embeddings of these drugs and their corresponding genes learned by X-Pert. As shown in Fig. 7B, ground-truth gene–drug pairs consistently showed higher cosine similarity than random drug–gene pairs for most genes across all three cell lines. These results indicate that X-Pert captures functional correspondence between genetic and chemical perturbations within a shared latent representation.

**Fig. 7.**
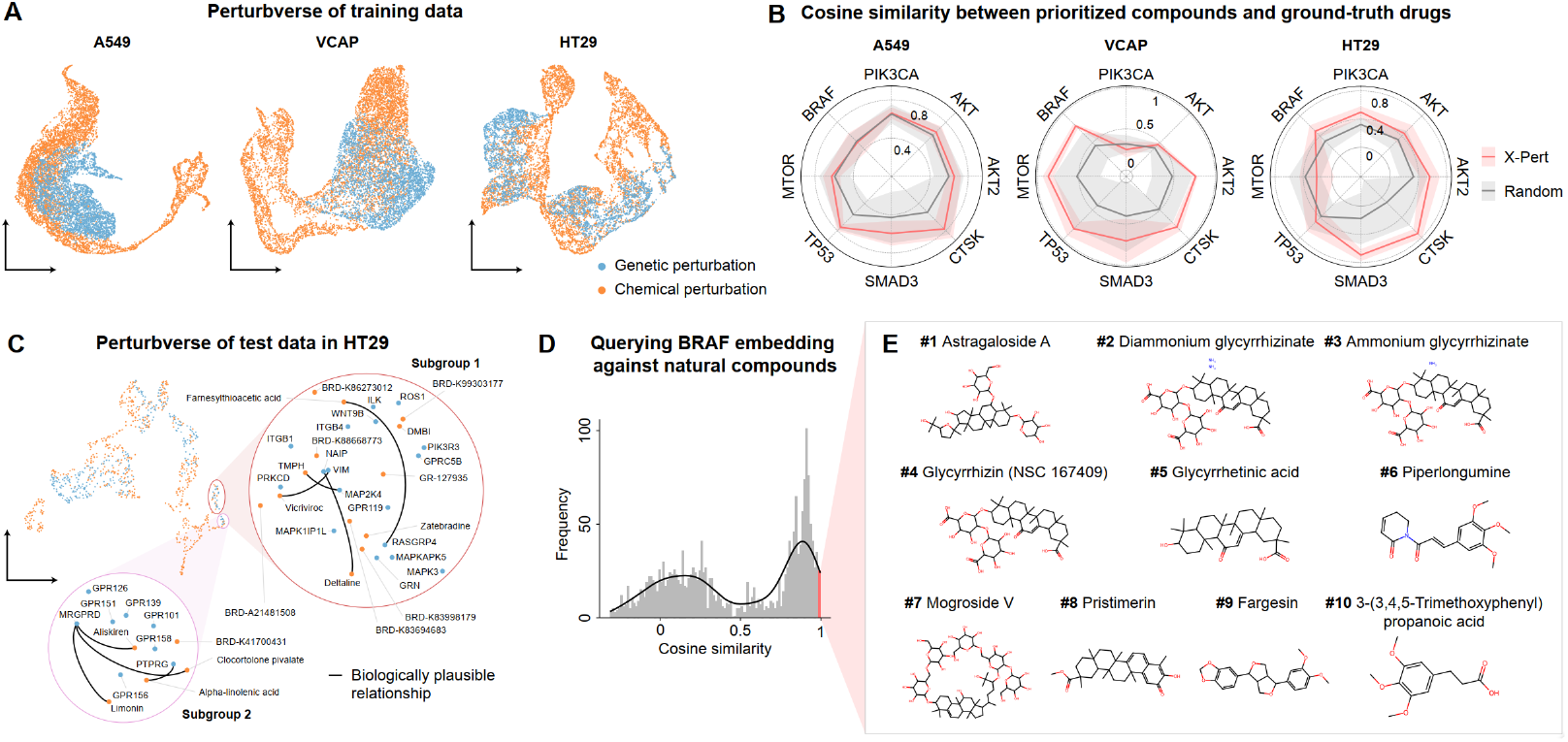
Cross-type alignment with genetic and chemical perturbations. **(A)** UMAP visualization of Perturbverse embeddings for all perturbations across A549, VCAP, and HT29 cell lines. **(B)** Radar plots comparing the cosine similarity between X-Pert–prioritized drugs and randomly selected drugs. Cosine similarity was calculated between the prioritized compounds and ground-truth drugs known to interact with the corresponding genes, as obtained from the DGIdb database. Lines denote the mean values, and shaded areas denote standard deviations across samples. **(C)** UMAP visualization of held-out test perturbations in the HT29 cell line, with the two highlighted subgroups shown with their corresponding perturbation names. Black lines denote biologically plausible relationships. **(D)** Histogram showing the distribution of cosine similarities between 1,919 natural compounds from the SelleckChem database and the BRAF perturbation embedding in the Perturbverse space. (E) Top ten drugs with the highest similarities with BRAF perturbation, shown with their rankings and compound names.

To assess its generalization, we focused on the HT29 cell line and examined the test perturbations that were excluded from model training (Fig. 7C). The low-dimensional visualization revealed that genetic and chemical perturbations were also well fused. Subgroups in this space revealed biologically plausible relationships. For example, in subgroup 1, deltaline aligned with NAIP deletion, both reducing muscle contraction despite distinct mechanisms^72, 73^; tetramethylpyrazine hydrochloride (TMPH) aligned with MAP2K4 knockout, both suppressing JNK signaling^74, 75^. Other examples included vicriviroc aligning with VIM knockdown, both of which impair tumor cell invasion^76, 77^, and farnesylthioacetic acid aligning with RASGRP4 knockdown, both suppressing Ras-dependent signal transduction and thus reducing proliferation and promoting apoptosis^78, 79^. Subgroup 2 showed parallels such as aliskiren with MRGPRD loss, both protecting cardiovascular cell function by attenuating angiotensin II (Ang II) signaling^80, 81^. Likewise, MRGPRD knockout showed functional similarity to anti-inflammatory drugs such as clocortolone pivalate and limonin^82-84^. Besides, both PTPRG loss and alpha-linolenic acid supplementation improved insulin sensitivity and reduced inflammation^85, 86^. Together, these results highlight X-Pert’s capacity to align genetic and chemical perturbations in a unified latent space and to uncover biologically meaningful relationships between perturbations.

We further evaluated X-Pert’s ability to perform cross-type perturbation retrieval within the Perturbverse of HT29 cells. HT29, a human colon cancer cell line harboring the BRAF V600E mutation, is frequently used to study the effectiveness of targeted therapies that inhibit the BRAF–MEK–ERK signaling pathway^87^. We queried the BRAF perturbation embedding of Perturbverse against 1,919 natural compounds from the Selleckchem database^88^, and we found a distinct subset of compounds with exceptionally high similarity scores to the BRAF perturbation embedding (Fig. 7D). We then identified top 10 candidates with high similarity (Fig. 7E). The top-one candidate is Astragaloside A, which, though not a direct enzymatic inhibitor of BRAF, have been shown to modulate the MAPK/ERK pathway^89^, thereby producing functional outcomes comparable to inhibition of RAF/MEK signaling. Another prominent group, including glycyrrhizic acid and its derivatives (including ammonium glycyrrhizinate and diammonium glycyrrhizinate), has been shown to act as RAS allosteric inhibitors^90^. We also identified Fargesin which has been reported to inhibit the PKA/CREB and p38 MAPK pathways, thereby affecting cellular processes regulated by the MAPK family^91^. While its mechanism does not directly target BRAF, its modulation of MAPK downstream kinases, such as ERK and p38^92^, suggests potential functional convergence with BRAF inhibition.

Taken together, these results demonstrate that X-Pert’s latent space can align genetic and chemical perturbations within a unified representation and support cross-type retrieval of compounds that retrieve compounds that recapitulate the effects of gene perturbations. This capability enables systematic translation between perturbation types, highlighting X-Pert’s potential as a general framework for integrative perturbation nalysis, drug discovery, and therapeutic repurposing.

## Discussion

In this study, we present X-Pert as a unified perturbation modeling framework that jointly represents perturbations and cellular states within a single architecture. By explicitly separating intervention encoding from cellular context modeling, X-Pert supports generalization across both perturbation types and cell types, moving perturbation modeling beyond collections of task-specific predictors toward a shared representation of intervention–response relationships conditioned on cellular context.

A central modeling insight underlying X-Pert is its multimodal formulation, which decouples intervention representation from cellular state representation. Perturbations are embedded independently of the cells on which they act, while cellular contexts are modeled through a context-aware encoder that captures intrinsic regulatory structure. This separation allows perturbation effects to be modeled as context-dependent transformations rather than fixed perturbation signatures, enabling generalization across unseen perturbations, combinatorial interventions, and diverse cell types within a single architecture. Biologically, this formulation reflects the fact that the same intervention can induce distinct responses in different cellular states, while similar transcriptional programs may arise from different interventions across contexts. The resulting Perturbverse provides a shared coordinate system that organizes perturbations by their induced transcriptional programs rather than experimental labels, enabling alignment between genetic and chemical interventions and facilitating translation from functional genomics to drug discovery.

The importance of such a general perturbation modeling framework becomes particularly evident under the current data regime of perturbation biology. As perturbation datasets increasingly span diverse intervention types, dosages, combinations, and cellular contexts, the primary challenge has shifted from data acquisition to organizing and integrating heterogeneous perturbations within a coherent modeling framework. Our results show that training on diverse perturbations and cell types improves predictive performance and generalizability, and demonstrated that X-Pert offers a practical way to accumulate, compare, and refine perturbation effects as new data are added. This behavior aligns naturally with the principles of foundation models, in which representation quality and generalization improve with increasing data diversity.

An important implication of general perturbation modeling is that the notion of intervention need not be restricted to molecular manipulations. While this study focuses on genetic and chemical perturbations, the underlying formulation treats perturbations abstractly as external operations applied to a cellular system. This abstraction naturally extends to non-molecular interventions, including changes in the cellular microenvironment, temperature, pH, nutrient availability, mechanical cues, or cell–cell interactions, all of which can induce systematic state transitions through shared regulatory mechanisms. By decoupling intervention representation from specific experimental implementations, frameworks such as X-Pert provide a conceptual foundation for integrating diverse forms of intervention within a single modeling paradigm.

Most existing perturbation models have so far focused primarily on transcriptomic profiles^22-27^, reflecting the widespread adoption of single-cell RNA sequencing (scRNA-seq) as the most mature and accessible technology for large-scale single-cell profiling. However, advances in sequencing and profiling technologies are rapidly expanding the omics modalities available for perturbation studies, including epigenomic readouts^93^ and protein expression measurements^94^ at single-cell resolution. These developments call for methods that can move beyond RNA-centric prediction to jointly capture multi-modal perturbation effects, encompassing RNA, DNA regulatory features, and protein-level responses. X-Pert’s modular design makes it readily extensible to additional omics layers, pointing toward perturbation foundation models that integrate multiple molecular modalities to capture the full spectrum of cellular responses to intervention.

A central challenge in advancing perturbation modeling lies in defining what it truly means for a model to be accurate. Conventional evaluation metrics, such as correlation across genes, often overestimate performance by rewarding the recovery of shared expression patterns rather than the reconstruction of perturbation-specific regulatory effects. This limitation is particularly consequential for general perturbation models, whose goal is to compare, transfer, and interpret effects across diverse interventions and cellular contexts. Our work emphasizes that evaluating predictive accuracy from the perspective of biological function is essential for assessing whether a model captures genuine regulatory responses rather than statistical regularities. Nevertheless, the metrics proposed here represent only an initial step, and future progress will require a broader suite of complementary evaluation strategies to disentangle causal perturbation effects from confounding factors and technical artifacts.

Finally, continued progress in perturbation modeling will depend not only on improved architectures but also on the availability of large, well-characterized perturbation corpora. Although our study paired original and perturbed cells, cellular responses remain inherently dynamic and noisy, and disentangling true perturbation signals from background variation remains challenging. By providing a general and extensible perturbation modeling framework, X-Pert offers a foundation upon which future efforts can build toward perturbation foundation models and, ultimately, predictive virtual cells capable of systematically reasoning about and engineering cellular responses across interventions and contexts.

## Methods

### Perturbation Perceiver

#### Perturbation Resampler

The Perturbation Perceiver, inspired by the perceiver architecture used in image–text multi-modal models^95^, is designed to integrate heterogeneous perturbations into a unified latent representation. Formally, let 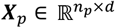 denote the perturbation embeddings, where *n*_*p*_ is the number of perturbation tokens and *d* is the latent dimensionality. We additionally introduce a set of learnable latent queries 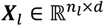, where *n*_*l*_ is fixed and independent of the type or number of perturbations. In the attention layer, the latent queries are used to extract information from both the perturbation inputs and the latent space itself. Specifically, the queries are defined as ***Q*** = ***X***_*l*_, while the keys and values are constructed by concatenating the perturbation embeddings with the latent queries: ***K*** = ***V*** = **[*X***_*p*_, ***X***_*l*_]. The attention operation is then given by

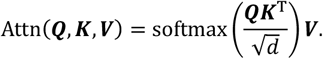

The output of the attention block is subsequently passed through a feed-forward network (FFN) with residual connections to enhance representational capacity and stabilize training:

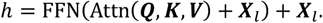

We set the number of latent queries *n*_*l*_ = 12 and embedding dimension *d* = 512. The Perturbation Perceiver consists of 2 attention layers, each employing a multi-head attention mechanism with 8 heads. The final output of Perturbation Perceiver is denoted as 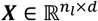, which serves as the keys and values for the downstream Cell Encoder. This design enables Perturbation Perceiver to project perturbations of diverse types into a shared latent space, providing a unified and flexible representation that can be seamlessly integrated by the Cell Encoder for subsequent modeling of perturbation–gene interactions.

### Perturbation-type–specific encoders

For different perturbation types, we employ type-specific encoders to transform the raw perturbation embeddings into the input space of the Perturbation Resampler. Specifically, for genetic perturbations we denote the original input as 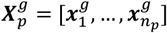, and for chemical perturbations we denote the input as 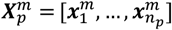, where 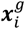 and 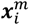 represent individual genetic and chemical perturbation embeddings, respectively. To ensure that perturbations of different types are mapped into a comparable latent space, we use separate three-layer multilayer perceptrons (MLPs) to project the original embeddings:

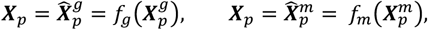

where *f*_*g*_ (⋅) and *f*_*m*_(⋅) denote the genetic and chemical encoders, respectively. These MLP-based encoders enable flexible handling of heterogeneous perturbation types while maintaining a unified representation that can be processed by the Perturbation Resampler.

For genetic perturbations, we leverage large language models (LLMs) to generate raw perturbation embeddings, motivated by the fact that LLMs capture broad biological knowledge across genes through large-scale pretraining. For each target gene, we constructed a natural language prompt of the form: “Impact of knocking out the gene {gene_name} on cell phenotype?” and submitted it to the GPT-4o model. The generated textual response, which encodes functional information about the gene perturbation, was then transformed into a numerical embedding using the GPT text-embedding-ada-002 model, resulting in a 1536-dimensional vector that serves as the raw genetic perturbation embedding.

For chemical perturbations, we adopted a cheminformatics-based approach. Each molecule was represented by its SMILES string, and the RDKit library^38^ was used to compute chemical embeddings via Morgan fingerprints. This procedure generated 1024-dimensional binary vectors, where each dimension (0/1) indicates the presence or absence of a specific chemical substructure feature.

Finally, to map perturbations from both types into a shared latent space, the perturbation-type–specific projection networks *f*_*g*_(⋅) and *f*_*m*_(⋅), were applied. These networks reduce the dimensionality of the raw embeddings from 1536 (genes) and 1024 (molecules) down to a unified 512-dimensional representation, which serves as the input for the Perturbation Resampler.

### Perturbation strength incorporation

The perturbation strength (dose or efficacy) can also be incorporated into the Perturbation Perceiver framework, encompassing both the dose information for chemical perturbations and the efficacy information for genetic perturbations. Formally, for input perturbation embeddings 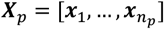, we denote the strength for each perturbation ***x***_*i*_ as *s*_*i*_ ∈ ℝ. To integrate this factor, we introduce a learnable vector ***w*** ∈ ℝ^*d*^ that modulates the embedding dimensions according to the perturbation strength. When strength information is included, the original perturbation embedding is modified as:

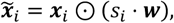

where ⊙ denotes element-wise multiplication. This formulation allows the perturbation embedding to be adaptively scaled according to its strength.

For genetic perturbations, the perturbation strength is equivalent to perturbation efficacy, which we estimate by comparing gene expression levels between original and perturbed cells. Let *μ*_*c*_ denote the average expression level of the target gene in original cells, and *μ*_*p*_ denote its expression level in perturbed cells. The perturbation efficacy is then defined as:

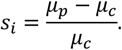

An efficacy value of −1 corresponds to complete knockout, meaning the target gene is reduced to zero expression relative to original cells.

For chemical perturbations, strength information is typically encoded as dose. For instance, in the Sciplex-3 dataset, each molecule is profiled under four concentrations: 10 nM, 100 nM, 1,000 nM, and 10,000 nM. We apply a log_10_ transformation to normalize the dose levels, yielding dose-based strength values that are more consistent for model integration: S_*i*_ = log_10_ dose.

In this way, the Perturbation Perceiver framework can incorporate quantitative measures of perturbation strength across types, providing a more nuanced and biologically informed representation of perturbations.

### Cell Encoder

#### Original cell input modelling

In the Cell Encoder, the input original cell is represented with three types of tokens: identity tokens, value tokens, and impact tokens. Following the design philosophy of previous single-cell foundation models, we treat the gene expression profile of a cell analogously to a sentence, where each gene corresponds to a word. Identity tokens are defined as gene identifiers, and we adopt the vocabulary from scGPT^25^ to represent them as the scGPT model supports efficient training. For genes not present in the original vocabulary, we extend the vocabulary by adding the new gene identifiers and initialize their embeddings using a uniform distribution matched to the mean and variance of the pretrained gene embeddings. For a given cell, the identity token sequence is thus represented as:

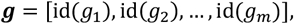

where id(*g*_*i*_) denotes the index of *g*_*i*_ in the vocabulary and *m* is the total number of genes.

The value tokens represent the expression levels of the corresponding genes and are denoted as

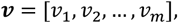

where *v*_*i*_ is typically the log-normalized expression value of gene *i*.

The impact tokens are designed to capture perturbation–gene interactions and vary depending on the perturbation type. For genetic perturbations, they are defined as

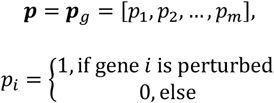

For chemical perturbations, they are defined as

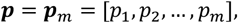

where *p*_*i*_ represents the affinity score between the protein encoded by gene *i* and the input molecule, calculated by inputting the protein sequence and the chemical SMILES string into a pretrained GraphDTA model.

The final embedding for an original cell is then obtained by summing the three projected components:

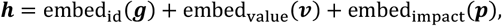

where embed_id_(⋅) projects gene identifiers into embeddings, embed_value_(⋅) is a nonlinear neural network mapping continuous expression values into embeddings, and embed_impact_(⋅) projects either binary perturbation indicators or affinity scores into embeddings depending on the perturbation type. All embedding dimensions are set to 512, consistent with the Perturbation Perceiver.

#### Transformer layers

The transformer layers in the Cell Encoder are designed to integrate the original cell input with perturbation information. Each layer is composed of two modules: a gated cross-attention layer that incorporates perturbation signals into the gene-expression representation, and a self-attention layer that models dependencies among genes.

In the gated cross-attention layer, cross-attention is performed between gene-expression tokens and perturbation embeddings, with an additional gating mechanism that adaptively regulates the strength of perturbation influence. Formally, given the perturbation embeddings ***X*** and the *l*-th layer output of gene-expression tokens ***H***^(*l*)^ ∈ ℝ^*m×d*^, the cross-attention is defined as:

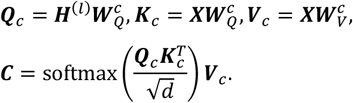

The gating mechanism is then applied to adaptively regulate the perturbation contribution:

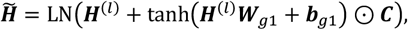

where LN(⋅) denotes layer normalization. The gated features are further refined with a gated feed-forward network:

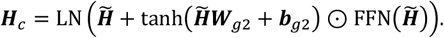

The output ***H***_*c*_ is then passed through a standard Transformer encoder block to capture gene–gene dependencies. Multi-head self-attention is applied as:

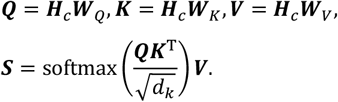

Residual connections and a feed-forward layer are applied to update the representation:

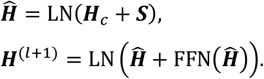

All ***W***_***\****_ and ***b***_***\****_ are trainable parameters. In summary, the entire encoder layer alternates between gated cross-attention and self-attention modules:

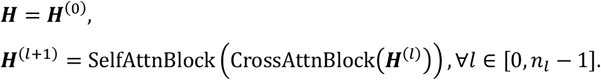

In this study, we set the number of encoder layers to *n*_*l*_ = 12, with 8 attention heads in each multi-head attention module.

#### Gene expression value decoder

The final output of the transformer layers is passed to the gene expression decoder, which predicts the complete gene expression profile of the perturbed cell. Let the output of the last transformer layer be denoted as 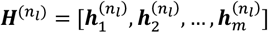, where 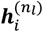 represents the latent embedding corresponding to gene *i*, and *m* is the total number of genes. For each output token, the decoder projects the latent representation into a scalar expression value:

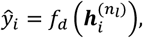

where *f*_*d*_(⋅) is a nonlinear projection layer implemented as a neural network.

Model training is performed by minimizing the mean squared error (MSE) loss between predicted and ground-truth perturbed expression values:

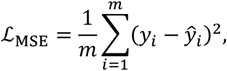

where *y*_i_ denotes the experimentally measured expression level of gene *i* under perturbation, and ŷ_*i*_ is the corresponding model prediction. This objective directly enforces accurate reconstruction of perturbation-induced transcriptional responses at the genome-wide level.

### Co-training across different perturbation types

For the X-Pert model trained jointly on genetic and chemical perturbations, we denote *k* as the total number of samples in a mini-batch, with *k*_*g*_ and *k*_*m*_ representing the number of genetic and chemical perturbation samples, respectively. To encourage alignment between perturbation types, we incorporate a Maximum Mean Discrepancy (MMD) loss that minimizes the distributional gap between the latent embeddings of gene perturbations and drug perturbations. The MMD loss is applied on the outputs of the perturbation-type– specific encoders, thereby enforcing the latent space to capture perturbation representations in a type-agnostic manner.

Formally, the MMD loss is defined as the squared distance between the kernel mean embeddings of the two distributions in a reproducing kernel Hilbert space (RKHS):

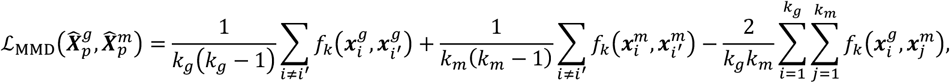

where f_k_(⋅,⋅) is the kernel function. We adopt the radial basis function (RBF) kernel, also known as the Gaussian kernel, defined as:

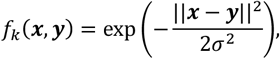

where ||***x*** − ***y***||^2^ denotes the squared Euclidean distance between vectors ***x*** and ***y***, and σ > 0 is the kernel bandwidth controlling the smoothness of the similarity measure. Following common practice, we set σ using the median heuristic, which has been shown to yield stable performance in high-dimensional spaces.

The overall training objective combines the MSE loss with the alignment loss:

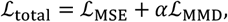

where *α* is a balancing hyperparameter. In this study, we set *α* = 0.2, which we found to provide a good trade-off between accurate reconstruction and effective cross–perturbation-type alignment.

#### Perturbverse embedding extraction

We use the output of the Perturbation Perceiver, 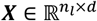, as the Perturbverse representation of the input perturbations. Since the number of latent queries *n*_*l*_ is fixed at 12, this design ensures that perturbations of any type, regardless of their original perturbation type or number of input tokens, are consistently projected into a standardized latent space of dimension 12×512. To obtain a compact perturbation embedding, the latent tokens are subsequently flattened into a 6,144-dimensional vector, which serves as the unified latent representation of the perturbation.

### Model evaluation

We used several metrics to perform the model evaluation and application.

### Pearson correlation coefficient (PCC)

The PCC measures the linear correlation between predicted (*ŷ*) and observed (*y*) values:

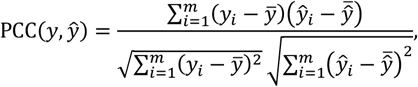

where 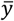 and 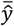 denote the means of the true and predicted values, respectively.

#### Coefficient of determination (*R*^2^)

The R^2^ score quantifies the proportion of variance in the observed data explained by the predictions:

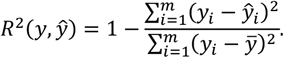

Higher values indicate stronger predictive power, with *R*^2^ = 1 representing perfect prediction.

#### Energy distance (E-distance)

The E-distance evaluates the dissimilarity between the distributions of perturbed and original cells, serving as a measure of perturbation effect size. Given two sets of samples ***X*** = **{*x***_1_, **…**, ***x***_*n*_**}** and ***Y*** = {***y***_1_, **…**, ***y***_*m*_**}**, the E-distance is defined as:

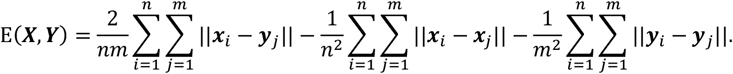

We use the scPerturb package to calculate the E-distance.

#### Cosine similarity

Cosine similarity measures the angle between two vectors and evaluates their directional similarity:

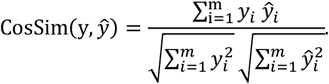

This metric is scale-invariant and emphasizes relative patterns rather than absolute magnitude.

#### Perturbation quantification with Hallmark gene sets

For each genetic perturbation, we evaluated pathway-level transcriptional changes by calculating gene set scores across all single cells. We used the Hallmark gene sets^45^ from the Molecular Signatures Database (MSigDB), which consist of 50 curated collections that capture and summarize specific, well-defined biological processes and cell states. These gene sets provide a compact and biologically interpretable representation of transcriptional programs, allowing us to assess whether perturbations induce consistent pathway-level responses. Gene set scores were computed using the Scanpy package^96^, specifically the ‘sc.tl.score_genes’ function, which calculates a per-cell enrichment score by comparing the expression of genes within each gene set against a reference background.

#### KNN accuracy

To quantitatively assess the discriminative ability of the learned Perturbverse embeddings (Fig. S2A), we employed a *K*-nearest neighbor (KNN) classification framework with stratified five-fold cross-validation. Specifically, a KNN classifier (*K* = 5) was trained on the PCA-transformed feature space, and the resulting predictions were evaluated using confusion matrices aggregated across folds. For each class (corresponding to a pathway label), a per-class accuracy was computed as the ratio of correctly classified samples to the total number of true samples in that class. This metric provides a fine-grained evaluation of how well the embedding preserves class-specific separability, independent of class imbalance.

### Databases and datasets

#### Candidate drug databases

We employed three different drug databases as candidate libraries to support perturbation retrieval and *in silico* drug screening tasks. For the drug perturbation retrieval experiment (Fig. 4G), we used the HERB 2.0 database^50^, which contains 30,916 natural compounds with curated annotations of chemical structures and pharmacological information. For the *in silico* drug screening task (Fig. 5D), we used an FDA-approved drug library from TargetMol^97^, consisting of 935 clinically approved compounds, ensuring the relevance of predictions to translational applications. Finally, for the cross-type perturbation retrieval analysis (Fig. 7E), we utilized a unique collection of 1,930 monomeric compounds derived from traditional Chinese medicine (TCM) curated by SelleckChem^88^, which provides structurally diverse natural products not typically represented in standard drug libraries.

#### Genetic and chemical perturbation data

We benchmarked X-Pert against multiple competing methods on genetic perturbation prediction using four datasets: Replogle2022_K562^7^, Replogle2022_RPE1^7^, and Norman2019^40^. The raw data of all the datasets were obtained from the scPerturb^19^ database.

For chemical perturbation benchmarking, we employed the Sciplex-3^14^ and CMAP^20^ datasets. The Sciplex-3 dataset is a single-cell perturbation dataset comprising 758,360 processed cells across three cell lines and 188 small-molecule perturbations after preprocessing. The CMAP dataset, which provides bulk-level gene expression profiles, contains 678,401 samples spanning 70 cell lines and 17,775 chemical perturbations after preprocessing.

All the datasets are processed with the same procedure following our previous benchmark^43^:

- Feature Selection: Top 5,000 highly variable genes (HVGs) were identified from each dataset using standard procedures. We also included all perturbed genes originally present in the dataset’s feature list.
- Normalization: Raw count data was normalized using the log-normalization method, implemented through the standard preprocessing workflow of Scanpy^96^.
- Original-cell pairing: For every cell subjected to perturbation, we randomly sampled an original cell from the same dataset to form a pre- and post-perturbation cell pair. These pairs served as the basis for training and evaluating models.
- Identifying differentially expressed genes (DEGs): For each perturbation, we identified DEGs by comparing perturbed cells to original cells. Genes were ranked based on adjusted *p*-values computed using the Wilcoxon test in Scanpy, with those having adjusted *p*-values < 0.05 designated as DEGs.

#### Tahoe-100M

Tahoe-100M is an exceptionally large-scale chemical perturbation dataset comprising approximately one million single cells. To reduce data redundancy and facilitate efficient training, we aggregated single-cell profiles into pseudo-bulk samples: for each combination for drug, cell line, and dose, the expression values of all corresponding single cells were averaged to yield a representative profile. After this preprocessing step, a total of 65,768 pseudo-bulk samples were retained for model training. To ensure that the model focuses on biologically informative signals, we further selected 10,000 highly variable genes (HVGs) across all samples as the feature set. This dimensionality reduction strategy retains the genes with the largest expression variability, thereby capturing the most relevant transcriptional changes induced by perturbations while reducing noise from less informative features.

#### *In silico* drug screening with X-Pert

To perform *in silico* drug screening, we combined disease-associated gene signatures with X-Pert–predicted gene expression profiles under drug perturbations. Disease signatures were obtained from the CRowd Extracted Expression of Differential Signatures (CREEDS) database^98^, which contains 839 annotated disease-versus-normal transcriptional signatures. For each disease, CREEDS provides two complementary gene sets: one consisting of up-regulated genes associated with the disease state, and another consisting of down-regulated genes. The rationale of our approach is that a drug with therapeutic potential should simultaneously suppress the disease-associated up-regulated genes while inducing the disease-associated down-regulated genes, thereby reversing the pathological transcriptional program.

We applied this framework to cell lines with defined disease associations in Tahoe-100M. For example, in the HepG2 cell line (originating from hepatocellular carcinoma, HCC), we used X-Pert to predict transcriptional responses to 831 candidate drugs across the 10,000 highly variable genes selected from the dataset. This produced drug-specific predicted gene expression profiles for HepG2 cells. The corresponding HCC gene signatures from CREEDS were then used to evaluate these predictions. Specifically, we computed two complementary enrichment scores using the Scanpy function ‘sc.tl.score_genes’: the up-regulated gene set score (up-score) and the down-regulated gene set score (down-score). Intuitively, lower up-scores and higher down-scores represent drugs with higher potential that could reverse disease signatures.

Large-scale chemical perturbation datasets such as Tahoe-100M are subject to technical batch effects, which in this case primarily arise from sequencing plate variability (Fig. 5B). These batch effects introduce systematic biases into X-Pert’s predictions, leading to spurious correlations between the calculated up-scores and down-scores. As a result, high up-scores often co-occur with proportionally high down-scores (Fig. 5D). This phenomenon is likely attributable to plate-specific biases in total transcript counts across different drug conditions, which can confound the biological signal and distort the prioritization of candidate drugs.

To mitigate these effects, we implemented a simple yet effective correction strategy based on linear regression. Specifically, we fit a regression line between the up-scores and down-scores across all candidate drugs for a given disease. The fitted line captures the dominant technical trend introduced by batch effects. We then identify drugs with the largest positive residuals (intercepts above the fitted line) as top candidates, since these drugs deviate most strongly from the batch-driven correlation and therefore likely reflect genuine biological activity rather than technical artifacts.

Drugs achieving large positive residuals (simultaneously low up-scores and high down-scores) are prioritized as candidate therapeutic compounds for the corresponding disease. This strategy allows us to systematically evaluate the potential of existing drugs to reverse disease-specific transcriptional dysregulation and highlights X-Pert’s utility in drug repurposing.

#### GDIdb drugs filtering

For the cross-type perturbation retrieval evaluation, we selected drugs known to interact with eight representative genes—AKT1, AKT2, CTSK, SMAD3, TP53, MTOR, BRAF, and PIK3CA—as the ground-truth references. For each gene, we obtained its interacting drugs from the DGIdb database^71^, which provides curated drug–gene interactions along with an interaction score that quantifies the strength of evidence supporting each relationship. To ensure reliability, we applied a threshold of 0.2 on the interaction score and retained only those drug–gene pairs exceeding this cutoff. After filtering, the number of drugs associated with each gene ranged from 7 to 23, forming the reference sets for evaluation.

## Supporting information

The supplemetary figures

## Data availability

The genetic perturbation data (Replogle2022_K562, Replogle2022_RPE1, and Norman2019) were downloaded from https://zenodo.org/records/10044268. The Sciplex-3 data were accessed from https://figshare.com/articles/dataset/sciplex3/22122701. The CMAP data was downloaded from Gene Expression Omnibus (GEO) with accession number GSE92742. The Tahoe-100M data were downloaded from https://huggingface.co/datasets/tahoebio/Tahoe-100M. The Nadig2025_HepG2 and Nadig2025_Jurkat data were acquired from GSE264667.

## Code availability

The code and documents of X-Pert are available at https://github.com/Chen-Li-17/X-Pert.

## Acknowledgements

The work is sponsored by The National Key R&D Program of China (2025YFC3409300), National Natural Science Foundation of China (62373210, 62433001, 92470105), and Tsinghua-Toyota Joint Research Institute Inter-Disciplinary Program (20243930093).

## Author contributions

C.L., L.W. and X.Z. conceived and designed the study. X.Z and L.W. supervised the study and provided critical revisions. C.L. collected the data, build the model, conducted the experiments, and analyzed the results. C.L. and L.W. drafted the manuscript and prepared the figures. X.Z. reviewed the manuscript and provided critical insights. All authors contributed to reviewing and approving the final manuscript.

## Reference

1. Kauffman, S.A. Metabolic stability and epigenesis in randomly constructed genetic nets. J Theor Biol 22, 437–467 (1969).

2. Jacob, F. & Monod, J. Genetic regulatory mechanisms in the synthesis of proteins. J Mol Biol 3, 318–356 (1961).

3. Linde, J., Schulze, S., Henkel, S.G. & Guthke, R. Data- and knowledge-based modeling of gene regulatory networks: an update. EXCLI J 14, 346–378 (2015).

4. Dixit, A. et al. Perturb-Seq: Dissecting Molecular Circuits with Scalable Single-Cell RNA Profiling of Pooled Genetic Screens. Cell 167, 1853–1866 e1817 (2016).

5. Adamson, B. et al. A Multiplexed Single-Cell CRISPR Screening Platform Enables Systematic Dissection of the Unfolded Protein Response. Cell 167, 1867–1882 e1821 (2016).

6. Datlinger, P. et al. Pooled CRISPR screening with single-cell transcriptome readout. Nat Methods 14, 297–301 (2017).

7. Replogle, J.M. et al. Mapping information-rich genotype-phenotype landscapes with genome-scale Perturb-seq. Cell 185, 2559–2575 e2528 (2022).

8. Rubin, A.J. et al. Coupled Single-Cell CRISPR Screening and Epigenomic Profiling Reveals Causal Gene Regulatory Networks. Cell 176, 361–376 e317 (2019).

9. Wessels, H.H. et al. Efficient combinatorial targeting of RNA transcripts in single cells with Cas13 RNA Perturb-seq. Nat Methods 20, 86–94 (2023).

10. Otto, J.E. et al. Structural and functional properties of mSWI/SNF chromatin remodeling complexes revealed through single-cell perturbation screens. Mol Cell 83, 1350–1367 e1357 (2023).

11. Aissa, A.F. et al. Single-cell transcriptional changes associated with drug tolerance and response to combination therapies in cancer. Nat Commun 12, 1628 (2021).

12. Chang, M.T. et al. Identifying transcriptional programs underlying cancer drug response with TraCe-seq. Nat Biotechnol 40, 86–93 (2022).

13. Zhao, W. et al. Deconvolution of cell type-specific drug responses in human tumor tissue with single-cell RNA-seq. Genome Med 13, 82 (2021).

14. Srivatsan, S.R. et al. Massively multiplex chemical transcriptomics at single-cell resolution. Science 367, 45–51 (2020).

15. Zhang, J. et al. Tahoe-100m: A giga-scale single-cell perturbation atlas for context-dependent gene function and cellular modeling. bioRxiv, 10.1101/2025.1102.1120.639398 (2025).

16. Papalexi, E. et al. Characterizing the molecular regulation of inhibitory immune checkpoints with multimodal single-cell screens. Nat Genet 53, 322–331 (2021).

17. Frangieh, C.J. et al. Multimodal pooled Perturb-CITE-seq screens in patient models define mechanisms of cancer immune evasion. Nat Genet 53, 332–341 (2021).

18. Yang, B. et al. PerturbDB for unraveling gene functions and regulatory networks. Nucleic Acids Res 53, D1120–D1131 (2025).

19. Peidli, S. et al. scPerturb: harmonized single-cell perturbation data. Nat Methods 21, 531–540 (2024).

20. Subramanian, A. et al. A Next Generation Connectivity Map: L1000 Platform and the First 1,000,000 Profiles. Cell 171, 1437–1452 e1417 (2017).

21. Huang, A.C. et al. X-Atlas/Orion: Genome-wide Perturb-seq Datasets via a Scalable Fix-Cryopreserve Platform for Training Dose-Dependent Biological Foundation Models. bioRxiv, 2025.2006. 2011.659105 (2025).

22. Roohani, Y., Huang, K. & Leskovec, J. Predicting transcriptional outcomes of novel multigene perturbations with GEARS. Nat Biotechnol 42, 927–935 (2024).

23. Bai, D., Ellington, C.N., Mo, S., Song, L. & Xing, E.P. AttentionPert: accurately modeling multiplexed genetic perturbations with multi-scale effects. Bioinformatics 40, i453–i461 (2024).

24. Hao, M. et al. Large-scale foundation model on single-cell transcriptomics. Nat Methods 21, 1481–1491 (2024).

25. Cui, H. et al. scGPT: toward building a foundation model for single-cell multi-omics using generative AI. Nat Methods 21, 1470–1480 (2024).

26. Schrod, S., Zacharias, H.U., Beissbarth, T., Hauschild, A.C. & Altenbuchinger, M. CODEX: COunterfactual Deep learning for the in silico EXploration of cancer cell line perturbations. Bioinformatics 40, i91–i99 (2024).

27. Yu, H., Qian, W., Song, Y. & Welch, J.D. PerturbNet predicts single-cell responses to unseen chemical and genetic perturbations. Mol Syst Biol 21, 960–982 (2025).

28. Xu, Y., Fleming, S., Tegtmeyer, M., McCarroll, S.A. & Babadi, M. Explainable modeling of single-cell perturbation data using attention and sparse dictionary learning. Cell Syst 16, 101245 (2025).

29. Hetzel, L. et al. Predicting Cellular Responses to Novel Drug Perturbations at a Single-Cell Resolution [C]. Proceedings of the 36th Conference on Neural Information Processing Systems (NeurIPS). 2022.

30. Piran, Z., Cohen, N., Hoshen, Y. & Nitzan, M. Disentanglement of single-cell data with biolord. Nat Biotechnol 42, 1678–1683 (2024).

31. Qi, X. et al. Predicting transcriptional responses to novel chemical perturbations using deep generative model for drug discovery. Nat Commun 15, 9256 (2024).

32. Maleki, S. et al. Efficient Fine-Tuning of Single-Cell Foundation Models Enables Zero-Shot Molecular Perturbation Prediction. arXiv preprint 2412.13478 (2024).

33. Klein, D. et al. CellFlow enables generative single-cell phenotype modeling with flow matching. bioRxiv, 10.1101/2025.1104.1111.648220 (2025).

34. Adduri, A.K. et al. Predicting cellular responses to perturbation across diverse contexts with STATE. bioRxiv, 10.1101/2025.1106.1126.661135 (2025).

35. Li, C. et al. TFcomb identifies transcription factor combinations for cellular reprogramming based on single-cell multiomics data. Genome Res 35, 1429–1439 (2025).

36. Roohani, Y.H. et al. Virtual Cell Challenge: Toward a Turing test for the virtual cell. Cell 188, 3370–3374 (2025).

37. OpenAI GPT-4 Technical Report. 2303.08774 (2023).

38. Bento, A.P. et al. An open source chemical structure curation pipeline using RDKit. J Cheminform 12, 51 (2020).

39. Nguyen, T. et al. GraphDTA: predicting drug-target binding affinity with graph neural networks. Bioinformatics 37, 1140–1147 (2021).

40. Norman, T.M. et al. Exploring genetic interaction manifolds constructed from rich single-cell phenotypes. Science 365, 786–793 (2019).

41. Ahlmann-Eltze, C., Huber, W. & Anders, S.J.N.M. Deep-learning-based gene perturbation effect prediction does not yet outperform simple linear baselines. Nat Methods, 1-5 (2025).

42. Kamimoto, K. et al. Dissecting cell identity via network inference and in silico gene perturbation. Nature 614, 742–751 (2023).

43. Li, C. et al. Benchmarking AI models for in silico gene perturbation of cells. bioRxiv, 10.1101/2024.1112.1120.629581 (2024).

44. Vinas Torne, R. et al. Systema: a framework for evaluating genetic perturbation response prediction beyond systematic variation. Nat Biotechnol (2025).

45. Liberzon, A. et al. The Molecular Signatures Database (MSigDB) hallmark gene set collection. Cell Syst 1, 417–425 (2015).

46. Radhakrishnan, S.K. et al. Transcription factor Nrf1 mediates the proteasome recovery pathway after proteasome inhibition in mammalian cells. Mol Cell 38, 17–28 (2010).

47. Oron, M. et al. The molecular network of the proteasome machinery inhibition response is orchestrated by HSP70, revealing vulnerabilities in cancer cells. Cell Rep 40, 111428 (2022).

48. Volmar, C.H. et al. M344 promotes nonamyloidogenic amyloid precursor protein processing while normalizing Alzheimer’s disease genes and improving memory. Proc Natl Acad Sci U S A 114, E9135–E9144 (2017).

49. Knox, C. et al. DrugBank 6.0: the DrugBank Knowledgebase for 2024. Nucleic Acids Res 52, D1265–D1275 (2024).

50. Gao, K. et al. HERB 2.0: an updated database integrating clinical and experimental evidence for traditional Chinese medicine. Nucleic Acids Res 53, D1404–D1414 (2025).

51. Lee, H.K., Oh, S.R., Kim, J.I., Kim, J.W. & Lee, C.O. Agastaquinone, a new cytotoxic diterpenoid quinone from Agastache rugosa. J Nat Prod 58, 1718–1721 (1995).

52. Mosca, L., Ilari, A., Fazi, F., Assaraf, Y.G. & Colotti, G. Taxanes in cancer treatment: Activity, chemoresistance and its overcoming. Drug Resist Updat 54, 100742 (2021).

53. Schneider, F., Samarin, K., Zanella, S. & Gaich, T. Total synthesis of the complex taxane diterpene canataxpropellane. Science 367, 676–681 (2020).

54. Gupta, A., Atanasov, A.G., Li, Y., Kumar, N. & Bishayee, A. Chlorogenic acid for cancer prevention and therapy: Current status on efficacy and mechanisms of action. Pharmacol Res 186, 106505 (2022).

55. Huang, S. et al. Chlorogenic acid effectively treats cancers through induction of cancer cell differentiation. Theranostics 9, 6745–6763 (2019).

56. Ibello, E. et al. Three-dimensional environment sensitizes pancreatic cancer cells to the anti-proliferative effect of budesonide by reprogramming energy metabolism. J Exp Clin Cancer Res 43, 165 (2024).

57. Yalniz, F.F. et al. A phase II study of addition of pracinostat to a hypomethylating agent in patients with myelodysplastic syndromes who have not responded to previous hypomethylating agent therapy. British journal of haematology 188, 404–412 (2020).

58. Sharman, J. et al. An open-label phase 2 trial of entospletinib (GS-9973), a selective spleen tyrosine kinase inhibitor, in chronic lymphocytic leukemia. Blood 125, 2336–2343 (2015).

59. Joshi, S. New insights into SYK targeting in solid tumors. Trends Pharmacol Sci 45, 904–918 (2024).

60. Rohila, D. et al. Syk Inhibition Reprograms Tumor-Associated Macrophages and Overcomes Gemcitabine-Induced Immunosuppression in Pancreatic Ductal Adenocarcinoma. Cancer Res 83, 2675–2689 (2023).

61. Varshosaz, J., Fard, M.M., Mirian, M. & Hassanzadeh, F. Targeted Nanoparticles for Co-delivery of 5-FU and Nitroxoline, a Cathepsin B Inhibitor, in HepG2 Cells of Hepatocellular Carcinoma. Anticancer Agents Med Chem 20, 346–358 (2020).

62. Choi, S.M. et al. Clioquinol, a Cu(II)/Zn(II) chelator, inhibits both ubiquitination and asparagine hydroxylation of hypoxia-inducible factor-1alpha, leading to expression of vascular endothelial growth factor and erythropoietin in normoxic cells. J Biol Chem 281, 34056–34063 (2006).

63. Huang, Z., Wang, L., Chen, L., Zhang, Y. & Shi, P. Induction of cell cycle arrest via the p21, p27-cyclin E,A/Cdk2 pathway in SMMC-7721 hepatoma cells by clioquinol. Acta Pharm 65, 463–471 (2015).

64. White, J.R., Jr. Empagliflozin, an SGLT2 inhibitor for the treatment of type 2 diabetes mellitus: a review of the evidence. Ann Pharmacother 49, 582–598 (2015).

65. Abdelhamid, A.M. et al. Empagliflozin adjunct with metformin for the inhibition of hepatocellular carcinoma progression: Emerging approach for new application. Biomed Pharmacother 145, 112455 (2022).

66. Scafoglio, C.R. et al. Sodium-glucose transporter 2 is a diagnostic and therapeutic target for early-stage lung adenocarcinoma. Sci Transl Med 10 (2018).

67. Luo, J., Hendryx, M. & Dong, Y. Sodium-glucose cotransporter 2 (SGLT2) inhibitors and non-small cell lung cancer survival. Br J Cancer 128, 1541–1547 (2023).

68. Lu, J. et al. Ciclopirox targets cellular bioenergetics and activates ER stress to induce apoptosis in non-small cell lung cancer cells. Cell Commun Signal 20, 37 (2022).

69. Ramalingam, S. et al. A Randomized, Double-Blind, Phase 2 Trial of Veliparib (ABT-888) With Carboplatin and Paclitaxel in Previously Untreated Metastatic or Advanced Non-Small Cell Lung Cancer. International Journal of Radiation Oncology+Biology+Physics 90, S4–S5 (2014).

70. Nadig, A. et al. Transcriptome-wide analysis of differential expression in perturbation atlases. Nat Genet 57, 1228–1237 (2025).

71. Cannon, M. et al. DGIdb 5.0: rebuilding the drug-gene interaction database for precision medicine and drug discovery platforms. Nucleic Acids Res 52, D1227–D1235 (2024).

72. Dobelis, P., Madl, J.E., Pfister, J.A., Manners, G.D. & Walrond, J.P. Effects of Delphinium alkaloids on neuromuscular transmission. J Pharmacol Exp Ther 291, 538–546 (1999).

73. Ahn, E.J. et al. Genotype-Phenotype Correlation of SMN1 and NAIP Deletions in Korean Patients with Spinal Muscular Atrophy. J Clin Neurol 13, 27–31 (2017).

74. Lin, J., Wang, Q., Zhou, S., Xu, S. & Yao, K. Tetramethylpyrazine: A review on its mechanisms and functions. Biomed Pharmacother 150, 113005 (2022).

75. Xue, Z. et al. MAP3K1 and MAP2K4 mutations are associated with sensitivity to MEK inhibitors in multiple cancer models. Cell Res 28, 719–729 (2018).

76. Velasco-Velazquez, M. et al. CCR5 antagonist blocks metastasis of basal breast cancer cells. Cancer Res 72, 3839–3850 (2012).

77. Berr, A.L. et al. Vimentin is required for tumor progression and metastasis in a mouse model of non-small cell lung cancer. Oncogene 42, 2074–2087 (2023).

78. Perez-Sala, D., Gilbert, B.A., Rando, R.R. & Canada, F.J. Analogs of farnesylcysteine induce apoptosis in HL-60 cells. FEBS Lett 426, 319–324 (1998).

79. Huang, S. et al. Ras guanine nucleotide-releasing protein-4 promotes renal inflammatory injury in type 2 diabetes mellitus. Metabolism 131, 155177 (2022).

80. Weng, L.Q. et al. Aliskiren ameliorates pressure overload-induced heart hypertrophy and fibrosis in mice. Acta Pharmacol Sin 35, 1005–1014 (2014).

81. Zhao, K. et al. Nuclear import of Mas-related G protein-coupled receptor member D induces pathological cardiac remodeling. Cell Commun Signal 21, 181 (2023).

82. Lan, L. et al. Mas-related G protein-coupled receptor D participates in inflammatory pain by promoting NF-kappaB activation through interaction with TAK1 and IKK complex. Cell Signal 76, 109813 (2020).

83. Song, C. et al. Limonin ameliorates dextran sulfate sodium-induced chronic colitis in mice by inhibiting PERK-ATF4-CHOP pathway of ER stress and NF-kappaB signaling. Int Immunopharmacol 90, 107161 (2021).

84. Scheinman, R.I., Gualberto, A., Jewell, C.M., Cidlowski, J.A. & Baldwin, A.S., Jr. Characterization of mechanisms involved in transrepression of NF-kappa B by activated glucocorticoid receptors. Mol Cell Biol 15, 943–953 (1995).

85. Brenachot, X. et al. Hepatic protein tyrosine phosphatase receptor gamma links obesity-induced inflammation to insulin resistance. Nat Commun 8, 1820 (2017).

86. Lin, J., Li, Q., Lei, X. & Zhao, H. The emerging roles of GPR158 in the regulation of the endocrine system. Front Cell Dev Biol 10, 1034348 (2022).

87. Chapdelaine, A.G., Ku, G.C., Sun, G. & Ayrapetov, M.K. The Targeted Degradation of BRAF V600E Reveals the Mechanisms of Resistance to BRAF-Targeted Treatments in Colorectal Cancer Cells. Cancers (Basel) 15 (2023).

88. Selleck products. https://www.selleckchem.com/ (2025).

89. Chen, T., Yang, P. & Jia, Y. Molecular mechanisms of astragaloside-IV in cancer therapy (Review). Int J Mol Med 47 (2021).

90. Zhang, Y. et al. Glycyrrhetinic acid binds to the conserved P-loop region and interferes with the interaction of RAS-effector proteins. Acta Pharm Sin B 9, 294–303 (2019).

91. Fu, T., Chai, B., Shi, Y., Dang, Y. & Ye, X. Fargesin inhibits melanin synthesis in murine malignant and immortalized melanocytes by regulating PKA/CREB and P38/MAPK signaling pathways. J Dermatol Sci 94, 213–219 (2019).

92. Shi, A., Liu, L., Li, S. & Qi, B. Natural products targeting the MAPK-signaling pathway in cancer: overview. J Cancer Res Clin Oncol 150, 6 (2024).

93. Metzner, E., Southard, K.M. & Norman, T.M. Multiome Perturb-seq unlocks scalable discovery of integrated perturbation effects on the transcriptome and epigenome. Cell Syst 16, 101161 (2025).

94. Saunders, R.A. et al. Perturb-Multimodal: A platform for pooled genetic screens with imaging and sequencing in intact mammalian tissue. Cell 188, 4790–4809 e4722 (2025).

95. Alayrac, J.B. et al. Flamingo: a Visual Language Model for Few-Shot Learning [C]. Proceedings of the 36th Conference on Neural Information Processing Systems (NeurIPS). 2022.

96. Wolf, F.A., Angerer, P. & Theis, F.J. SCANPY: large-scale single-cell gene expression data analysis. Genome Biol 19, 15 (2018).

97. TargetMol. https://www.targetmol.com (2025).

98. Wang, Z. et al. Extraction and analysis of signatures from the Gene Expression Omnibus by the crowd. Nat Commun 7, 12846 (2016).

