## Supplementary material for "Unified multimodal learning enables generalized cellular response prediction to diverse perturbations": The supplemetary figures

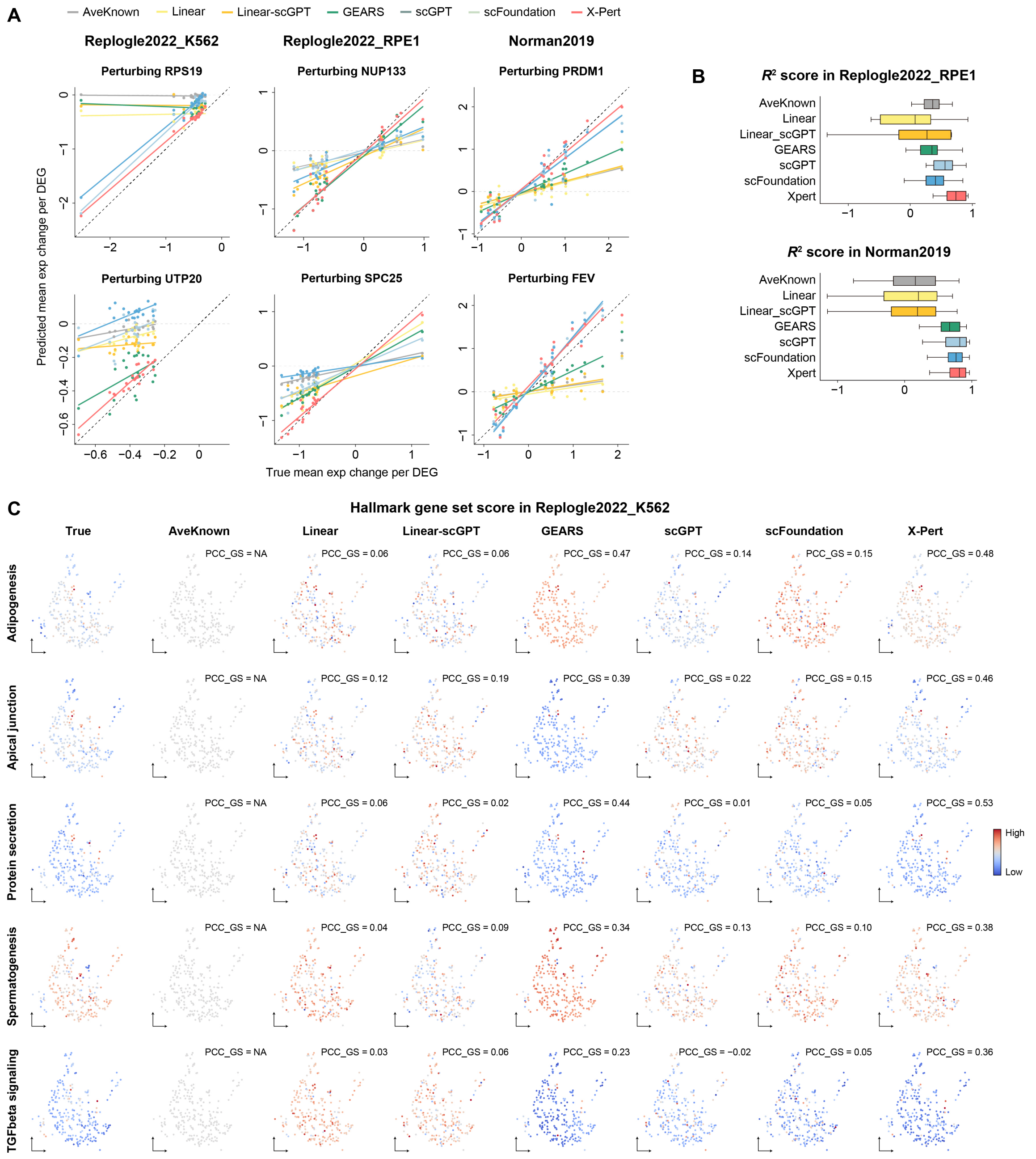


**Supplementary Fig. 1. (A)** Comparison between predicted and ground-truth expression changes of the top 20 DEGs in three datasets. Dots represent the mean predicted expression changes from each method, and a linear regression fit was added to visualize the correlation between the predicted and true mean values. For each dataset, two representative genetic perturbation are shown on each row. **(B)** Box plots comparing the $R^2$ score between predicted and observed top 20 DEGs in Replogle2022_RPE1 and Norman2019. **(C)** UMAP visualization comparison of gene set scores on several representative Hallmark gene sets.

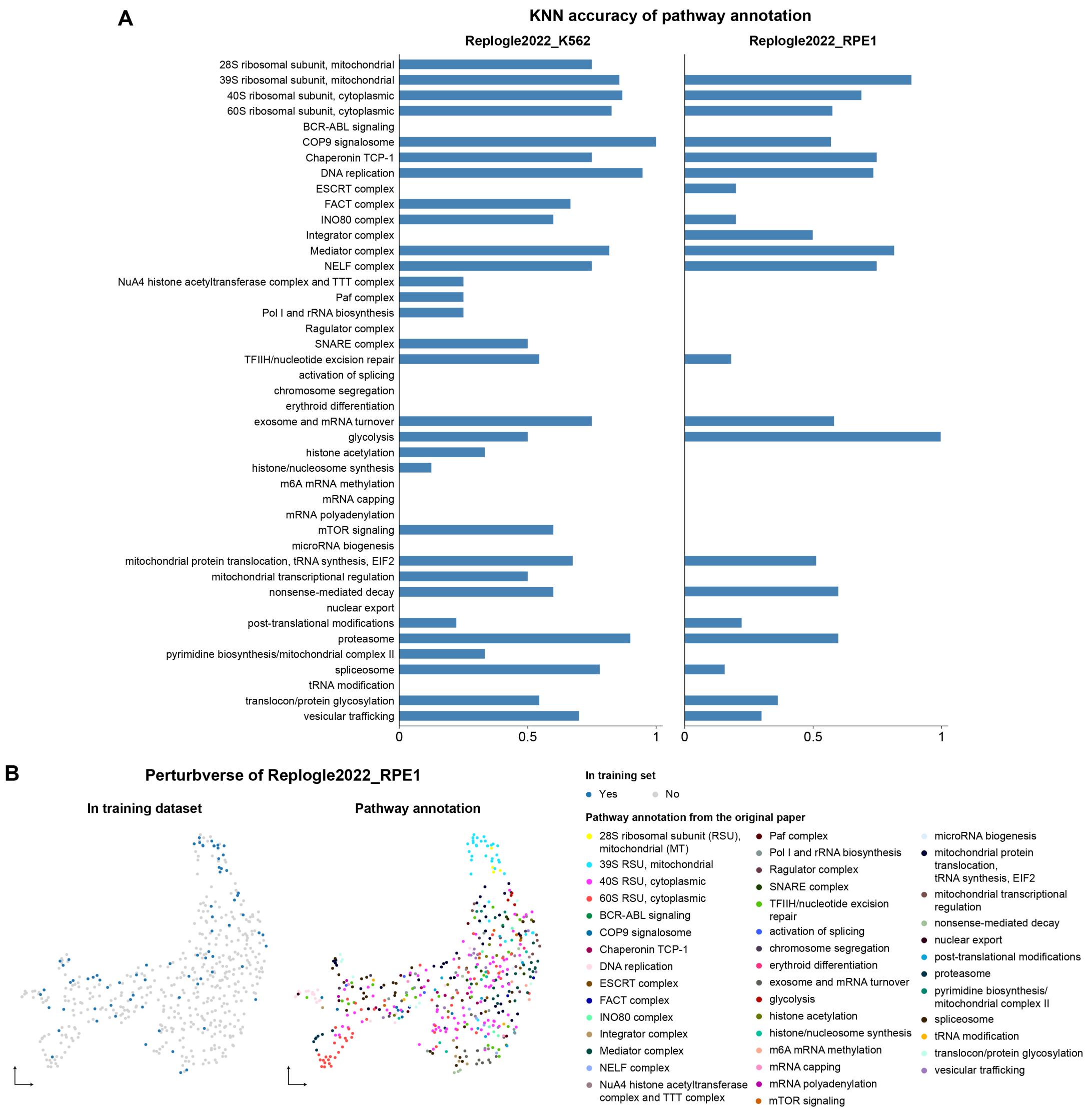


**Supplementary Fig. 2. (A)** Bar plots showing the KNN classification accuracy of latent genetic perturbation embeddings across pathway annotations manually curated from the original dataset in Replogle2022_K562 and Replogle2022_RPE1, respectively. (B) UMAP visualization of 504 genetic perturbations from Replogle2022_RPE1, with pathway annotations manually curated from the original dataset.

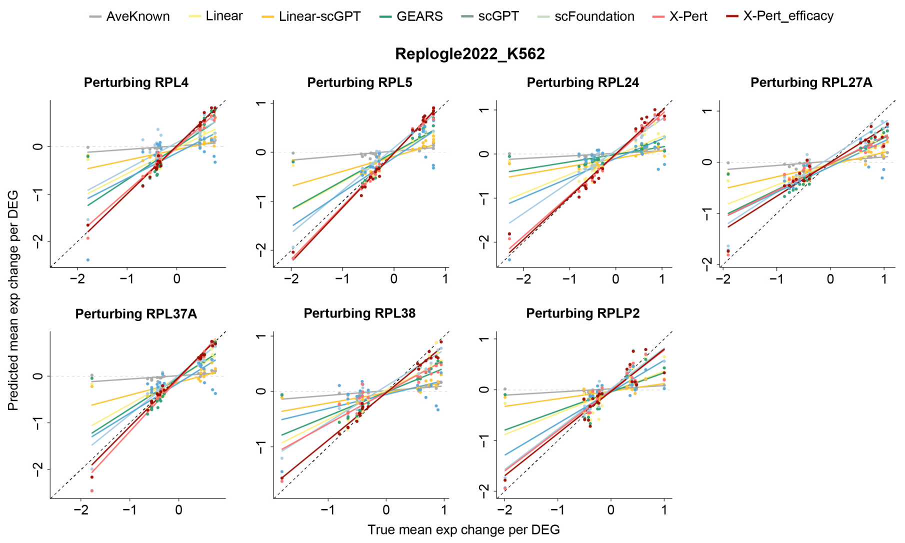


**Supplementary Fig. 3.** Comparison between predicted and ground-truth expression changes of the top 20 DEGs in the Replogle2022_K562 dataset. Dots represent the mean predicted expression changes from each method, and a linear regression fit was added to visualize the correlation between the predicted and true mean values. These perturbations are RPL-related test perturbations.

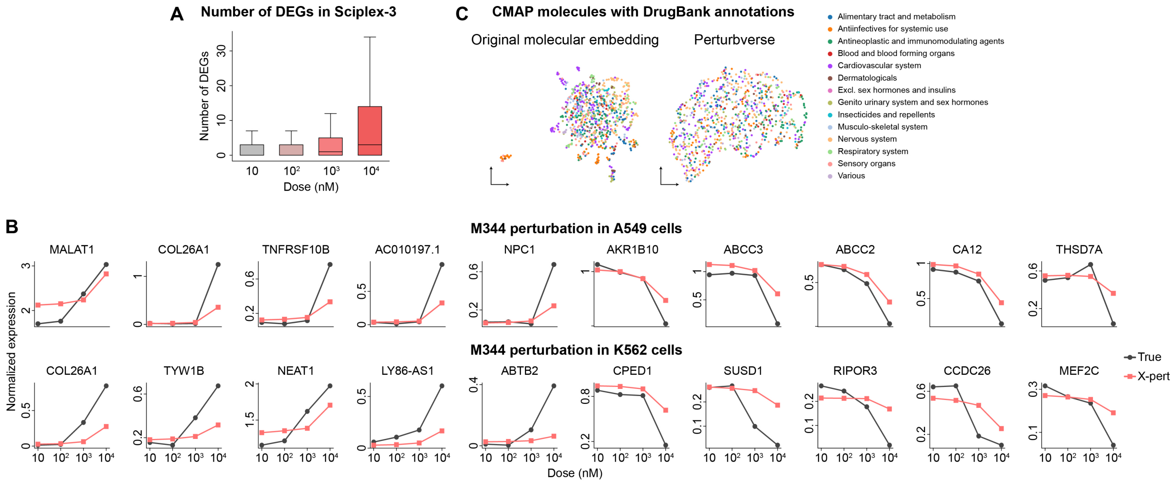


**Supplementary Fig. 4.** (A) Number of DEGs in the Sciplex-3 dataset. (B) Line plots showing X-Pert–predicted gene expression levels versus ground-truth measurements across four molecular doses. The top and bottom rows correspond to the drug M344 in cell lines A549 and K562, respectively. (C) UMAP visualization of chemical perturbations in the original molecular embedding space and the Perturbverse, colored by ATC Level 1 annotations.
